# PRDM9-mediated meiotic hotspot specification is constrained in humans despite extensive sequence diversity

**DOI:** 10.64898/2026.09.10.750150

**Authors:** Rachel L. Cosby, Marja Brolinson King, Alexandra M. Poch, M. Blake Evans, Dawn E. Watkins-Chow, Briana Young, Sherry Ralls, Veronica Gomez-Lobo, Kenneth I. Aston, Donald F. Conrad, Todd S. Macfarlan

## Abstract

PRDM9 specifies meiotic recombination hotspots through a rapidly evolving C2H2 zinc-finger (ZNF) coding minisatellite that determines DNA-binding specificity. Although this minisatellite harbors extraordinary allelic diversity in humans, the functional consequences of most naturally occurring variants remain unknown. Here we functionally characterize 80 human PRDM9 alleles using genome-wide chromatin profiling. Despite extensive sequence diversity within the ZNF array, most alleles function indistinguishably from common A and C hotspot-specifying alleles, revealing that human PRDM9 function is more constrained than its sequence variation predicts. In contrast, rare variants identified in infertile individuals occupy two functional extremes: abundant and novel DNA binding specificity or minimal DNA binding, suggesting that both gain- and loss-of-function alleles may disrupt symmetric hotspot specification during meiosis, plausibly contributing to human infertility. Together, our findings define the functional landscape of human PRDM9 variation and provide a framework for interpreting the impact of newly discovered PRDM9 alleles.

## Introduction

Although infertility affects 1 in 6 people worldwide^1^, unexplained infertility accounts for ∼33% of cases^2^ and poses a significant economic and health burden^3^. One cause of infertility is failed implantation or spontaneous miscarriage due to chromosome aneuploidies (trisomy or monosomy)^4^. During normal gametogenesis, diploid cells replicate their DNA and divide via two rounds of meiosis to generate haploid gametes that retain exactly one copy of each chromosome^5^. To ensure that chromosomes are segregated properly, meiotic recombination (MR) physically links the homologous chromosomes together^5^. Defects in MR increase with age and resulting aneuploidies are associated with quantitative sperm defects^6,7^ and account for an estimated 10% of pregnancy losses^4^. Thus, variants in MR regulators may represent unexplored genetic risk factors for infertility.

During prophase I of meiosis, chromosomes condense and MR is initiated by SPO11, which generates double-strand DNA breaks (DSBs) at sites throughout the genome^5^. Following end resection, the cleaved strand invades the homologous chromosome, resulting in break repair via either a non-crossover or crossover event, the latter of which links the chromosomes together. In either case, DNA is exchanged between the homologous chromosomes, shuffling genetic material between homologs and contributing to genome evolution^5^. While the major steps of MR are conserved in almost all sexually reproducing eukaryotes, the DSB sites and the proteins involved vary across species^8,9^. In some species, programmed DSBs occur more frequently at specific genomic sites, called “hotspots”^8^. Hotspots in species such as yeast, plants, and birds, are enriched near promoter regions and are denoted by tri-methylation of K4 on the histone H3 tail (H3K4me3)^8^. In many vertebrates, including humans and most mammals, hotspots occur away from promoters at regions specified by PRDM9, a Krüppel-associated box containing zinc finger protein (KRAB-ZFP)^10–13^.

PRDM9 consists of several protein domains, including a KRAB domain and PR/SET methyltransferase domain at the N-terminus and a tandem array of C2H2 zinc fingers (ZNFs) at the C-terminus. PRDM9 specifies hotspot locations by binding to genomic DNA via its ZNF array and trimethylating both K4 and K36 residues on the tails of adjacent histone H3 proteins (H3K4me3/H3K36me3 dual mark)^14,15^. Consistent with its important role in MR, *Prdm9^-/-^* mice are sterile, with hotspots reverting to ancestral promoter regions^16,17^. Coding and non-coding heterozygous PRDM9 variants have been associated with both non-obstructive azoospermia (NOA, lack of sperm)^18–20^ and primary ovarian insufficiency (POI)^21,22^, suggesting PRDM9 is haploinsufficient. Although the mechanism underlying PRDM9-mediated infertility is not fully understood, PRDM9 binding to the same ∼1kb region on both homologs (symmetric) is associated with beneficial meiotic outcomes, including increased crossover rate^23,24^, synapsis^25,26^, and homolog engagement^27^ as well as improved DNA break repair^24,25^. Conversely, PRDM9 binding to only one homolog (asymmetric) is associated with negative fertility outcomes and inter-subspecies sterility^28^.

One prediction of a model favoring symmetric binding is that some PRDM9 variants, especially those with distinct binding motifs, might be incompatible due to reduced symmetric binding in meiosis. This is likely to happen for two reasons: 1) PRDM9 is a fast-evolving gene, with over 100 alleles in humans and mice and dozens in other species^10,29–42^ and 2) most sequence variation in PRDM9 occurs in its DNA binding domain, either in the number of ZNFs or in the specific amino acids, called “fingerprint residues,” within each ZNF (positions -1, +3, and +6) that determine DNA binding specificity^43^. While the cause of this rapid evolution is unknown, the sequence encoding *PRDM9*’s ZNF array is highly repetitive and prone to errors during DNA replication, consistent with its high mutation rate in germline and somatic tissues^31^. Additionally, computational modeling studies^44,45^ suggest that the rapid evolution can be partially explained by “erosion” of PRDM9 binding sites in the genome. This occurs when PRDM9 binds more strongly to one of the two homologs, leading to preferential DSB formation on the “hotter” allele that is then repaired by the “colder” allele^13^. PRDM9 therefore removes its own binding motifs from the genome, putting selective pressure on novel PRDM9 variants to restore symmetric binding^44,45^ and effective MR.

If PRDM9 variants have sufficiently different binding specificity, such as in the case of interspecies hybrids^46,47^, loss of symmetric binding could result in reduced fertility due to meiotic arrest or increased aneuploidies. Recent studies that have identified heterozygous truncation and missense PRDM9 variants in individuals with either NOA^18–20^ or POI^21,22^, support this assertion, as PRDM9 haploinsufficiency would be predicted to reduce symmetric binding^44,45^. Because most minor human PRDM9 alleles occur in the heterozygous state with a “major” (ie: high frequency) allele^29^, it is plausible that some extant PRDM9 variants might impair fertility if their binding patterns are sufficiently distinct from the major allele. The functional consequences of PRDM9 allelic divergence on hotspot specification are unclear, however, because standard methods to profile DNA binding (ie: ChIPseq) are challenging for PRDM9 and are not scalable to tens or hundreds of variants, leading to data being available for only a few human (A and C) and mouse (Dom2 and Cst) variants^11,48–51^. Additionally, while algorithms predicting ZNF binding based solely on AA sequence have utility, they often fail to correctly predict empirically determined DNA binding motifs^52,53^. To address these limitations, we developed a scalable, high-throughput assay to determine the genome-wide binding properties of PRDM9 variants, which we then applied to 80 human variants. In doing so, we characterized each variant’s molecular properties and found that most naturally occurring human PRDM9 alleles are functionally indistinguishable from the major A or C alleles. In contrast, PRDM9 variants identified in infertile individuals are functional outliers that likely decrease binding symmetry when paired with the major A allele.

## Results

### Classification of human PRDM9 variants

We first surveyed previous studies^10,12,29–31,54,55^ to collect the sequences of known PRDM9 variants (n=87, Table S1). As previously reported^29–33^, most human PRDM9 variation occurs either in the number of zinc fingers in the tandem array or in the fingerprint residues, suggesting these variants may specify different hotspot locations. Previous studies analyzing motif enrichment at known human recombination hotspots ^11,50^ and PRDM9 binding data ^48,49^ have demonstrated that the C-terminal zinc fingers (PRDM9-A ZNFs 6-10; PRDM9-C ZNFs 6-11) determine DNA binding specificity, whereas the N-terminal fingers are thought to be structural or involved in PRDM9 multimerization^48,49,56^. Considering only the fingerprint residues, we manually aligned each variant in a stepwise fashion, prioritizing shared fingerprint combinations and fingerprints that differed by a single amino acid (see Methods).

Consistent with previous research^29^, we found two broad categories of variants, which we refer to as A-like (n=65) or C-like alleles (n=22) (Fig. 1, Supplemental Table 1). A-like and C-like variants differ primarily in their C-terminal fingerprint patterns (A: NHR-DHR-DNS-NHR-NHR and C: NHR-DHS-DNS-WVR-DNS-NHR). Almost all PRDM9 variants include additional, degenerate ZNFs upstream (WHI) and downstream (DSY) of the array. A-like variants are on average shorter than C-like variants (mean # total ZNF = 11.8 vs 13.5 respectively, *p =3.9e^-5^,* Wilcoxon rank sum test, W=307, df=1), due to having fewer C-terminal DNA binding ZNFs (mean # DBD ZNF = 4.9 vs 6.1, *p =7.5e^-9^,* Wilcoxon rank sum test, W=165.5, df = 1; Fig. 1). For both A-like and C-like variants, several are predicted to specify identical hotspots as their cognate alleles due to either identical fingerprint residues (n=12 for A, n=1 for C) or identical C-terminal fingerprints (n=22 for A, n=9 for C). Most human PRDM9 variation occurs in the N-terminal ZNFs, which would not be expected to impact PRDM9 DNA binding specificity.

**Figure 1:**
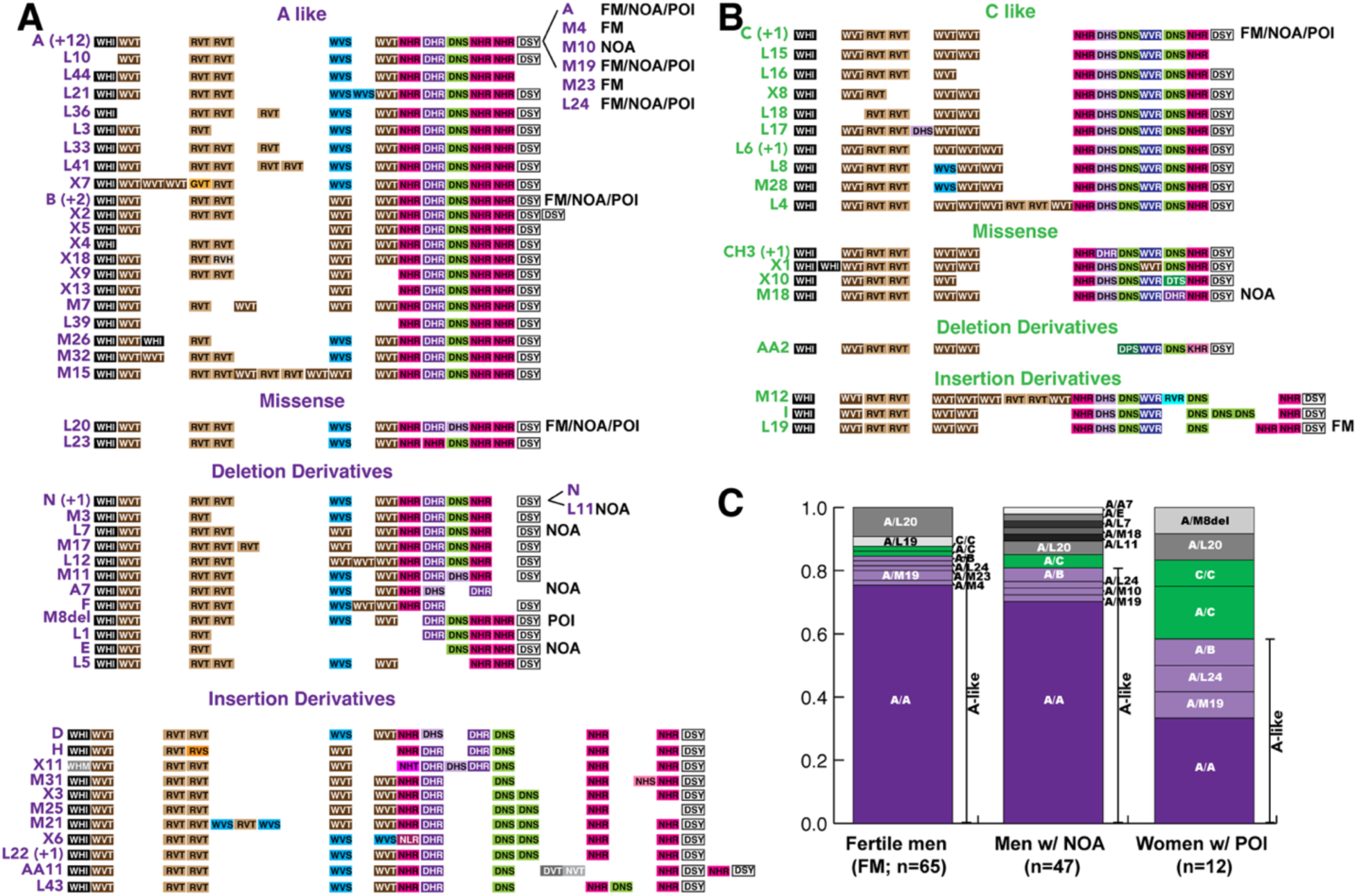
Extensive human PRDM9 allelic diversity includes both highly similar and distinct alleles, including novel alleles identified in infertile individuals. A-B) Visual representations of the ZNF fingerprint residues for previously described and novel human PRDM9 variants, categorized as either A-like (A; purple) or C-like (B; green) based on C-terminal fingerprint residues. Bright colors represent ZNFs predicted to contact DNA; neutral colors represent predicted multimerization ZNFs. Numbers in parentheses represent additional PRDM9 variants with identical ZNF array fingerprints. C) Bar chart describing frequency of diploid genotypes in in cohorts of fertile men (FM), men with non-obstructive azoospermia (NOA), and women with primary ovarian insufficiency (POI). Genotypes with A-like variants alone are shaded in purple, genotypes with at least one C like variant are shaded in green, and missense, insertion, or deletion derivatives of either A or C are shaded gray. Two novel variants were identified in infertile patients (A7 in NOA and M8del in POI). Variants identified in either of the three cohorts are noted with labels in A-B.

Despite the similarities, a subset of PRDM9 variants differ from the A/C-like alleles (Fig. 1, Supplemental Table 1). Some alleles contain one or more additional ZNFs within the C-terminal region (“insertion derivatives”, n=12 for A-like, n=3 for C-like), while others contain fewer ZNFs (“deletion derivatives”, n=13 for A-like, n=1 for C-like). There are also variants within each group with C-terminal fingerprint missense mutations (n=2 for A-like, n=5 for C-like). Given that the C-terminal region is responsible for binding DNA, it is possible that these divergent variants specify unique hotspots relative to the A/C alleles.

### Some infertile individuals possess rare PRDM9 variants

Because previously reported PRDM9 alleles were identified in individuals of unknown fertility status, we identified candidate fertility-impairing PRDM9 variants using long-read sequencing of *PRDM9*’s ZNF array^29^ (see Methods) in cohorts of healthy control men (n=65), men with NOA (n=47), and women with POI (n=12) (Supplemental Table 2). Most individuals in both male cohorts were homozygous for the A/A variant (n=49, 75.4% control; n=33, 70.2% NOA; Fig. 1C, Supplemental Table 2), consistent with the high frequency of the A allele in Caucasian populations^29^. Of the remaining control males, six possessed variants with identical ZNF 6-10 fingerprint residues as A (A/B, n=1; A/L24, n=1; A/M19, n=2; A/M23, n=1; A/M4, n=1) and another six with an A-like minor variant (A/L20), all of which would be predicted to bind either identical or highly similar sites as the A allele. We also observed a small number of genotypes with C-like variants (A/C, n = 1; C/C, n = 1; and A/L19, n = 1). While several of these allele combinations were also observed in men with NOA (A/B, n=2; A/L24, n=1; A/M19, n=1; A/L20, n=2; A/C, n =2), we identified several genotypes (n=5; 10.6%) unique to the NOA cohort, including a C-like missense variant (M18) and four A-like deletion-derivatives (E, A7, L11, L7), all heterozygous with the A allele (Fig. 1C, Supplemental Table 2). Of these, one variant is unique to this study (A7) and two are rare variants (E = 0-3% and M18 = 0-1% allele frequency respectively^29^). We also observed several minor alleles (n=8, 77%) in our POI cohort, including a novel A-like deletion-derivative allele (M8del) (Fig. 1C, Supplemental Table 2). While an increased prevalence of rare variants in NOA and POI patients might be due to demographic differences between our relatively small cohorts (Supplemental Table 2), it motivated us to assess their molecular functionality in comparison to the canonical A/C alleles.

### PRDM9-mediated H3K4me3 deposition recapitulates in vivo PRDM9 binding

To assess the molecular capabilities of PRDM9 variants, we sought to determine their genome-wide DNA binding properties. To overcome technical limitations in directly assessing PRDM9 binding and to improve scalability, we adapted previously described approaches that inferred PRDM9 binding by identifying sites of new H3K4me3 deposition in the presence of PRDM9 (ie: PRDM9-dependent H3K4me3)^49,57,58^. We first transfected HEK293T cells, which lack endogenous PRDM9 expression, with a construct (Supplemental Data 1^59^) encoding either GFP or a PRDM9 variant. We then used Cleavage Under Targets and Release Using Nuclease (CUT&RUN)^60^ to identify sites of PRDM9-dependent H3K4me3 as a proxy for PRDM9 binding (Fig. 2A).

**Figure 2:**
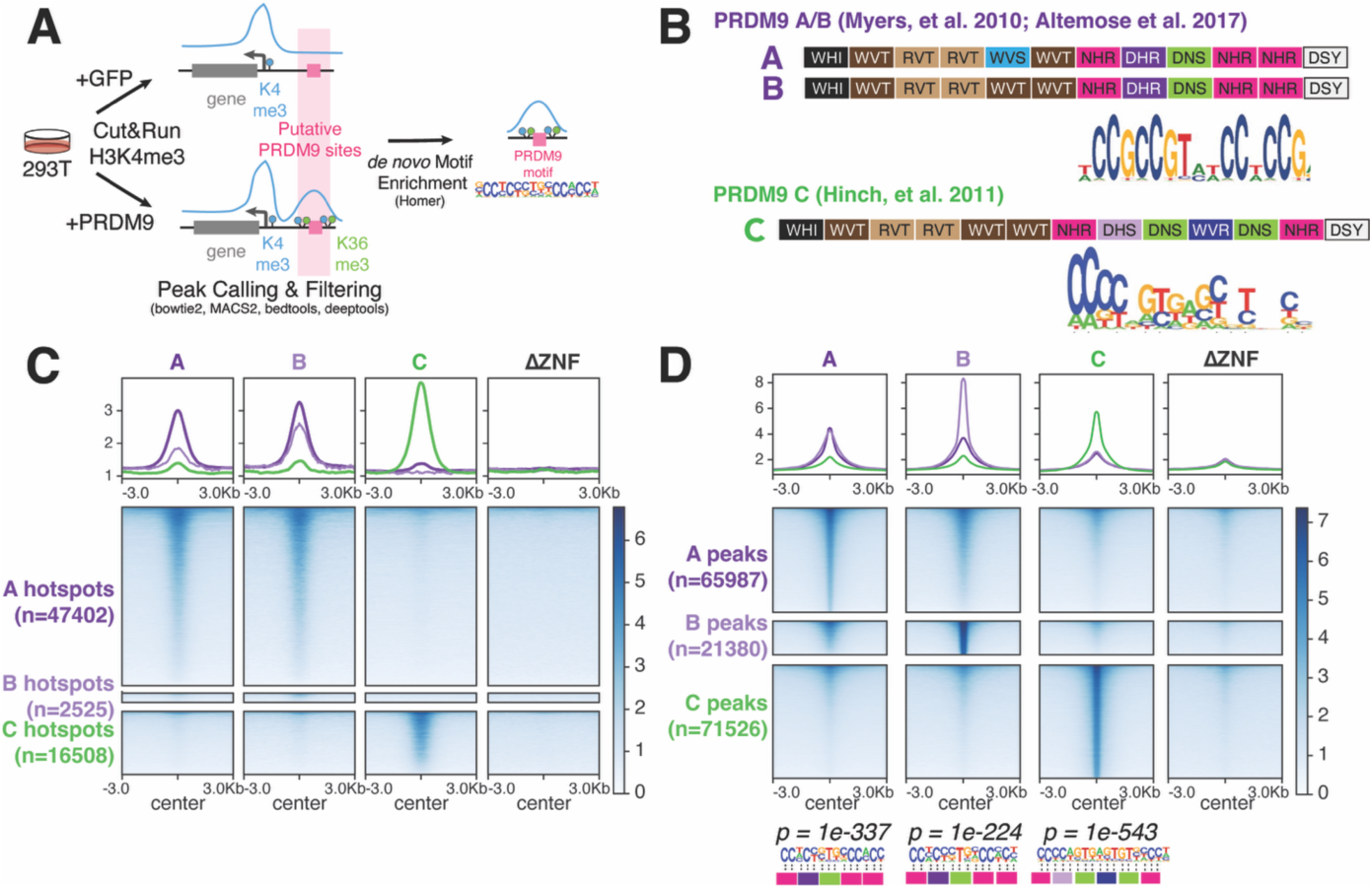
A high throughput CUT&RUN assay recapitulates features of in vivo PRDM9 binding. A) A CUT&RUN based assay to determine PRDM9 binding using H3K4me3 as a proxy. Cells are transfected with vectors encoding either GFP or a PRDM9 variant, and H3K4me3 sites are assessed genome-wide in the presence of these proteins. Predicted PRDM9 binding sites are those that gain H3K4me3 in the presence of PRDM9 but not GFP. B) ZNF array fingerprint residues and empirically determined motifs for the A, B, and C variants^11,33,48^, with motifs represented as a sequence logo. C) Heatmaps plotting mean CUT&RUN H3K4me3 signal at previously identified A, B, and C meiotic hotspots^50^ in the presence of PRDM9 A, B, or C. Results are contrasted with a ZNF-less PRDM9 (ΔZNF) as a negative control. D) Heatmaps plotting mean CUT&RUN H3K4me3 signal at A, B, and C peaks, with logos showing the most enriched motif sequence within each variant’s respective peaks shown below.

To verify that our assay reflects PRDM9 binding *in vivo,* we first benchmarked using variants whose binding patterns and motifs are known (A/B/C)^11,29,33,50,61^. We assessed PRDM9-dependent H3K4me3 deposition for the A (*n=4*), B (*n=1*), and C (*n=2*) variants, with a PRDM9 variant engineered to lack the zinc-finger array (ΔZNF; *n=1*) as a negative control, at recombination hotspots previously identified in A/A, A/B, and A/C individuals^48,50^ (Supplemental Table 3). As expected, we found that A and B variants, which are known to bind identical hotspots and motifs (Fig. 2B), deposited H3K4me3 at A and B sites, but not C sites, and vice versa for the C allele (Fig. 2C). Despite its intact SET domain, PRDM9-ΔZNF did not deposit H3K4me3 at any known recombination hotspots (Fig. 2C). These dissimilarities were not due to PRDM9 variant expression differences, as all variants (A, B, and C) were expressed at comparable levels in HEK293T cells (Extended Data 1).

We then identified putative PRDM9 binding sites for each PRDM9 variant by calling H3K4me3 peaks (MACS2^62^), filtered to remove promoters and H3K4me3 peaks present in matched GFP samples (n=7; bedtools^63^), and merged replicate peak sets (Extended Data 2) where applicable to generate a master list of peaks identified in two or more samples (bedtools^63^). Consistent with previous studies^48,49^, we found that canonical PRDM9 variants methylate many genomic sites (A: *n=65987*; B: *n=21380*; C: *n=71526*), with minimal overlap between A/B and C variants and almost no methylation for PRDM9ΔZNF (Fig. 2D). Further, A/B peaks and C peaks were enriched for known A ([A] *p=1e-04*, Z=337.07; [B] *p=1e-04,* Z=404.14) or C (*p=1e-04;* Z=364.88) hotspots (regioneR^64^, empirical *p,* n=10000 shuffles)^50^. We then identified enriched motifs within bound regions (homer^65^) for each variant and recovered the known motifs (Fig. 2B, Supplemental Data 2^59^) as the top hit for each peak set (A *p = 1e-337*, B *p = 1e-224*, and C *p = 1e-543;* binomial distribution; Fig. 2D).

To test how our approach compared to previous studies, we performed analogous experiments assessing PRDM9-dependent H3K4me3 using ChIPseq as a readout (Supplemental Table 3). While there were some differences between the two approaches, such as increased replicate variability for CUT&RUN samples (Extended Data 2), the overall conclusions from the ChIPseq experiments were the same: PRDM9-A (n=2) and PRDM9-C (n=2) deposit H3K4me3 at their known hotspot sites (Extended Data 3), and PRDM9-dependent H3K4me3 peaks were enriched for their respective hotspots (A: *p=1e-04,* Z=612.42; C: *p=1e-04*, Z=336.78; regioneR^64^, empirical *p,* n=10000 shuffles) and motifs (A *p=1e-1031;* C *p=1e-1087;* binomial distribution; Extended Data 3, Supplemental Data 2^59^). We observed similar results when we directly assessed PRDM9 binding (anti-FLAG ChIP) under the same conditions (FLAG-PRDM9 A or C, n=2 each) and found that all three datasets (direct PRDM9 binding [ChIP] and PRDM9-dependent H3K4me3 [CUT&RUN and ChIP]) were highly concordant (Extended Data 3-4; Supplemental Table 3). Collectively, these results demonstrate that PRDM9-dependent H3K4me3 deposition is an effective proxy for PRDM9 binding and that our CUT&RUN-based assay recapitulates key features of PRDM9 binding *in vivo* comparably to existing approaches.

### Most human PRDM9 variants bind canonical motifs at A or C hotspots

We then utilized our assay to identify the DNA binding properties, including genomic binding sites (peaks) and motifs, for previously uncharacterized PRDM9 variants (n=75 CUT&RUN, n=3 ChIP; Supplemental Tables 3-4). We observed a wide range in binding capability for PRDM9 variants, with the median A-like and C-like variants binding 16604 and 56092 genomic sites respectively (Fig. 3A). A-like variants bound significantly fewer sites than C-like variants (*p=0.014,* Wilcoxon rank-sum test, W = 394, df =1). Both A (n=10682) and C (n=16982) bind fewer sites than the median variant of their type, suggesting their binding motifs have begun to erode. Several variants bound substantially fewer (<15%: A-like < 4738, C-like < 16982) or more (>85%: A-like > 73563, C-like > 137642) sites compared to other variants of their type (Fig. 3A), suggesting differences in binding motif frequency genome-wide or differences in DNA binding capability.

**Figure 3:**
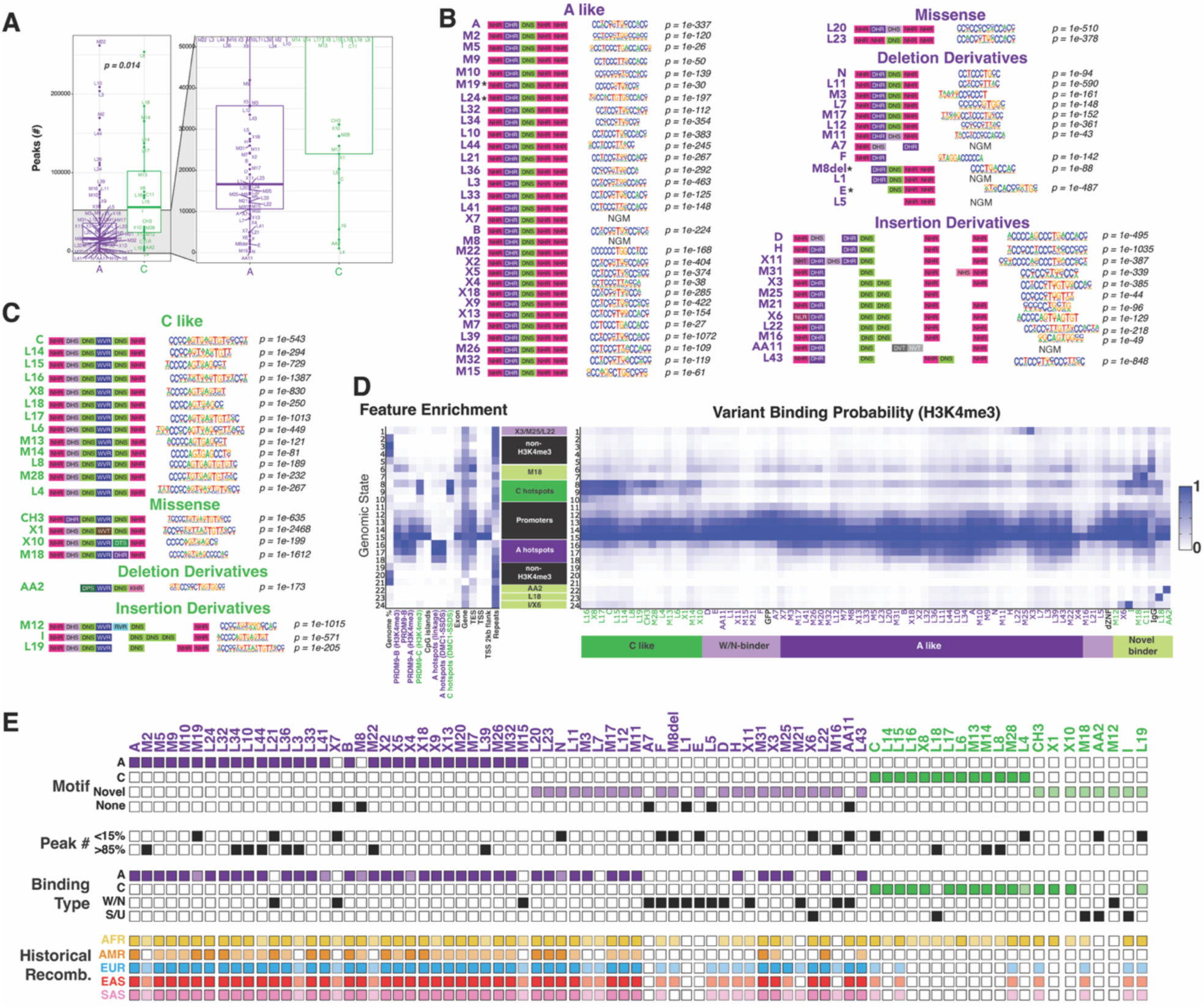
Human PRDM9 variants include canonical, strong, and weak DNA binders. A) Boxplot summarizing the number of binding peaks identified for each tested PRDM9 variant, colored and grouped according to type (A or C). Inset shows data range from 0 – 50,000 peaks. C variants on average bind significantly more sites than A variants (*p = 0.014*, Wilcoxon rank-sum test, W = 394, df =1). B-C) DNA-contacting ZNF fingerprint diagrams paired with the top enriched motif within peaks for each variant. *p* values derived from the binomial distribution. NGM = no good motif was identified (see Methods). Variants whose motifs were determined via ChIPseq are denoted with a star. D) 24-state ChromHMM model incorporating all CUT&RUN H3K4me3 data. Left panel shows column-wise fold enrichment of different genomic features for each state (blue = maximum value, white = minimum value). Right panel shows the binding probability (emission probabilities) for each variant across the 24 states. Middle panel shows states annotated using information from both feature enrichment and variant emission probabilities. Binding classes for PRDM9 variants (canonical A-like or C-like, weak/non-binder [W/N], and strong/unique [S/U] binder) are shown below. E) Summary of binding properties derived from H3K4me3 CUT&RUN/ChIPseq data, including motif type (dark purple = A, dark green = C, light purple = novel A-like, light green = novel C-like, black = NGM), variants binding either <15% or >85% sites genome wide, binding type (canonical A [dark purple]/C[dark green], weak A [light purple]/C [light green], W/N, or S/U), and mean historical recombination rates across super-populations assessed in the 1000 Genomes Project (dark box = mean per generation recombination rate *r* > 1e-8, light box = mean *r* > 2e-9, empty box = no evidence of historical recombination). AFR = African, AMR = American, EUR = European, EAS = East Asian, SAS = Southeast Asian.

Using our motif analysis pipeline (homer^65^), we found that variants of both types (A/C) with identical C-terminal fingerprint residues, with few exceptions, bound identical motifs (Fig. 3B-C, Supplemental Data 2^59^), consistent with our ZNF-based prediction. Similarly, variants with one or more residue changes in their C-terminal fingerprints relative to the corresponding major allele (A: L20 and L23; C: X1, X10, M18, and CH3) bound near identical motifs, differing only by a single DNA triplet (Fig. 3B-C, Supplemental Data 2^66^). These results suggest most variants likely bind the same genomic sites as either the A or C alleles respectively.

To assess the binding patterns of PRDM9 variants, we modeled the PRDM9-dependent H3K4me3 status (CUT&RUN) genome-wide using ChromHMM^67^ (see Methods). Using a 24-state model (Fig. 3D, Supplemental Table 5, Supplemental Data 3^59^), we identified states that represented promoter (State 15 [S15]; CpG island: 48.25 fold enriched [FE], transcription start site [TSS]: 43.3 FE) and promoter-adjacent regions (mean ± SD) (S11-14; CpG island: FE = 7.01 ± 7.58, TSS: 5.55 ± 5.53 FE), non-H3K4me3 regions (S2-5 and S19-21; *P(H3K4me3)* = 0.04 ± 0.036), and states enriched for known A (S16-18; FE = 28.95 ± 9.5) and C (S8-10; FE = 19.38 ± 12.45) hotspots^29,50^ (Fig. 3D, Supplemental Table 5, Supplemental Data 3^59^). As expected, H3K4me3 from all samples, including PRDM9 variants and GFP controls, was found at promoters (S15; *P(H3K4me3)* = 0.98 ± 0.06) (Fig. 3D; Supplemental Tables 3,5). Outside of these regions, PRDM9 A (n=39) and C (n=15) like variants with either identical or highly similar arrays generally bound exclusively to their cognate hotspots (A: *P(H3K4me3:S16-18) =* 0.43 ± 0.23 vs *P(H3K4me3:others) =* 0.07 ± 0.08; C: n= *P(H3K4me3:S8-10)* = 0.40 ± 0.23 vs *P(H3K4me3:others) =* 0.08 ± 0.09). Thus, most PRDM9 alleles tested (54/72) likely specify identical hotspots as the major A/C alleles. However, a subset of A/C-like alleles (A: L41, N, X4, and X7; C: L4, L19), hereafter referred to as “weak canonical” binders, bound comparatively fewer peaks (< 15%) than other variants of their type, suggesting there may be differences in binding affinity even amongst highly similar alleles (Fig. 3 A, E). Collectively, these results demonstrate that extensive PRDM9 allelic diversity does not necessarily result in functional diversity.

### Highly variable PRDM9 alelles likely specify unique hotspots

Although most PRDM9 alleles appear to be functionally redundant, 24% (18/75) have distinct binding patterns and motifs compared to the major alleles and fall into two broad classes: “strong/unique” and “weak/non” binders (Fig. 3E; Supplemental Table 4). We considered variants to be strong/unique binders if they had unique binding motifs, high H3K4me3 deposition outside of known promoter or hotspot regions, and/or bound more genomic sites (>85^th^ percentile) than other alleles of the same type. With one exception (X6), these were all C-like variants (M18, L18, AA2, I), and all bound unique genomic states relative to other PRDM9 variants (Fig. 3D). To infer the cause of these differences, we inspected the protein sequence of the ZNF array for each strong/unique variant. In the case of AA2 (*P(H3K4me3:S22)* = 0.80), substantial differences in its DBD ZNFs (ZNF=DPS-WVR-DNS-NHR) relative to the C allele (ZNF=NHR-DHS-DNS-WVR-DNS-NHR) results in a shorter motif and fewer binding sites (n=3148 peaks). In contrast, M18 differs from the C allele by only one ZNF finger (ZNF11 = DHR vs DNS) and binds a motif that is otherwise C-like except for a change from GTG to CCC. This change leads to abundant (71760 peaks) and unique binding (*P(H3K4me3:S6-7)* = 0.58 ± 0.19 vs *P(H3K4me3:non-promoters)* = 0.17 ± 0.20) genome wide (Fig. 3D). L18 (*P(H3K4me3:S23)* = 0.74) has an identical DBD ZNF fingerprint pattern (ZNF=NHR-DHS-DNS-WVR-DNS-NHR) as C but its shorter motif suggests that it only uses a subset of its DBD ZNF array (ZNF=NHR-DHS-DNS-WVR-NHR) (Fig. 3B-C). While the reason for this is unclear, it is possible that a missense mutation (E299D) in the linker region between fingers DNS and NHR might affect L18’s binding specificity. I (*P(H3K4me3:S24)* = 0.82) also has a shorter binding motif, despite having a longer DBD ZNF array (ZNF=NHR-DHS-DNS-WVR-DNS-DNS-DNS-NHR) than C, suggesting additional ZNFs do not always increase the length of the binding motif (Fig. 3B-D). Although X6 has an A-like DBD array (ZNF=NLR-DHR-DNS-DNS-NHR-NHR), its binding most resembled the I variant (*P(H3K4me3:S24)* = 0.31), including an identical binding motif (Fig. 3B-C). Collectively, these results suggest that a variety of ZNF array changes can lead to novel PRDM9 binding.

We classified variants as weak/non binders if they met one or both of the following criteria: 1) low H3K4me3 deposition outside of promoter regions (*P(H3K4me3:non-promoters)* = 0.06 ± 0.08) comparable to control samples (ie: GFP/PRDM9ΔZNF; *P(H3K4me3:non-promoters)* = 0.1 ± 0.09), and/or 2) no identifiable binding motif (Fig. 3B-C). These criteria gave us a total of 12 weak/non binders, all of which were A-like (A7, AA11, D, E, F, M8del, L1, L21, L5, M15, M16, M21, X11) except for M12 (C-like). Some variants (L21, M15, M21) bind an A-like motif, but only weakly bind A regions (mean *P(H3K4me3:S16-18 =* 0.12 ± 0.09; Fig. 3C). A-insertion derivatives D, X11, and M16 have unique binding motifs, but unlike strong/unique binders appear to bind diffusely genome-wide (*P(H3K4me3:non-promoters)* = 0.07 ± 0.08). F also binds uniquely but only to a small fraction of sites genome wide (n=2671 peaks). Other variants with no identifiable motif (AA11, A7, L1, and L5), may not bind DNA at all (Fig. 3B). Notably, all but two (L21 and M15) are either insertion or deletion derivatives. While M15 has a canonical A-like DBD array, its multimer ZNFs (multiZNFs) is four ZNFs longer than the major A variant (Fig. 1A). This is also true of other variants with weak binding, including X7 and L41 (+2 multiZNFs) and L4 (+4 multiZNFs) (Fig. 1A). These results suggest there may be an optimal range of ZNF array lengths for proper PRDM9 function, and that ZNF changes outside the DBD portion of the array can affect binding. Collectively, these results imply that most weak/non-binder variants specify substantially fewer meiotic hotspots compared to canonical alleles. Importantly, these differences are likely not due to PRDM9 expression levels, as most variants are comparably expressed in 293T cells (Extended Data 1) and expression level overall does not track with observed binding patterns (ie: weak/nonbinder variants are not expressed at lower levels than canonical variants, and vice-versa for strong/unique variants).

### Historical recombination rate provides insight into PRDM9 variant age and functionality

To better understand the relative age and potential functionality of PRDM9 variants, we leveraged previously published fine-scale historical recombination data for 26 populations, representing 5 super-populations (Africa, Southeast Asians, East Asians, Europeans, and Americans), from the 1000 Genomes Project^68,69^. Specifically, we plotted the mean per-generation recombination rate (*r*) for each population over all peak regions for each PRDM9 variant (Supplemental Data 4^59^; see Methods). If a variant’s binding sites showed evidence of recombination over background (PRDM9ΔZNF peaks, mean *r* = 2e-9) for a given population we infer that the hotspots specified by the variant have been utilized within that population, whereas an absence of historical recombination at peak loci suggests either that the variant has recently emerged or does not specify functional hotspots. As before, we first benchmarked this method using the well characterized A/C variants. We found elevated recombination at PRDM9-A peak loci for all populations (mean *r* = 1e-8 [American/African] to 8e-8 [European/East Asian/Southeast Asian]) (Fig. 3E and Extended Data 5; Supplemental Table 4, Supplemental Data 4^59^). For the C variant, we observed strong evidence of recombination in African populations (mean *r* = 6e-9 to 1.2e-8) and marginal recombination in European and Asian populations (mean *r* = 4e-9 to 6e-9) but no recombination in American populations (mean *r* < 2e-9) (Fig. 3E and Extended Data 5; Supplemental Table 4, Supplemental Data 4^59^). These patterns are broadly consistent with the known allele frequencies of the A/C alleles in human populations^10,12,29–31,54,55^.

These trends were identical for other canonical A- and C-like variants (Fig. 3D and Extended Data 5; Supplemental Table 4, Supplemental Data 4^59^), with a few exceptions. Three canonical A-like variants (M2, L3, and M22), which bind to a much larger number of sites than other A variants (>85%), show reduced recombination rates in all populations (mean *r* = 3e-9 – 6e-9), suggesting they may specify additional novel or low-frequency hotspots. While nearly all C-like variant binding sites show evidence of recombination in African populations, many lack recombination in other populations, suggesting they are likely at low frequency in non-Africans.

For the noncanonical variants, which include the strong/unique and weak/non binders, we observe two broad patterns. Most strong/unique binders (except AA2) had at least some evidence of recombination at bound sites, though less than the canonical variants (Fig. 3E and Extended Data 5; Supplemental Table 4, Supplemental Data 4^59^). This, combined with the large number of peaks and novel motifs, suggest that these variants are functional but comparatively young. This may also be true for a subset of weak/non binders (M15, F, M8del, L5, D, X11, M21, and M16), suggesting they retain some functionality but are either younger or less functional than canonical alleles (Fig. 3E and Extended Data 5; Supplemental Table 4, Supplemental Data 4^59^). In contrast, the recombination rate at peaks bound by some weak/non-binder variants (A7, L1, E) are indistinguishable from sites bound by our negative control (PRDM9ΔZNF), which, in conjunction with minimal evidence of genomic binding in our assay, suggests that these variants may not be functional (Fig. 3E and Extended Data 5; Supplemental Table 4, Supplemental Data 5^59^). Notably, this includes two variants (A7, E) that were identified in our azoospermia cohort, supporting the possibility that these variants may contribute to patient infertility.

### PRDM9 allelic dominance influences putative hotspot locations

Our data indicates that some PRDM9 alleles have different binding properties compared to the canonical A and C alleles. Importantly, most alleles occur in the heterozygous state with A or C^29^, and previous studies have shown hotspot specification in heterozygous individuals reflects a mixture of the PRDM9 variants involved, with one variant contributing more (ie: “dominant”) than the other^29,49,50^. Although the mechanistic basis for this dominance is not entirely understood, it seems to be a function of two intersecting phenomena: 1) PRDM9 molecules multimerize via their ZNF arrays^49,56^ and 2) the binding properties of the multimer are affected by the frequency of each variant’s binding sites in the genome, with “dominant” alleles being those whose binding sites have not eroded^29,49,50^. To better understand the functional consequences of different PRDM9 variants, we adapted our CUT&RUN assay to mimic heterozygous individuals by co-expressing two PRDM9 variants, prioritizing high frequency (>5% genotype frequency in at least one population^29^) genotypes (A/C, A/B, A/L4, A/L14, A/L19, A/N) as well as combinations including one major allele and either a strong/unique (A/M18, A/I) or weak canonical or weak/nonbinder allele (A/A7, A/E, C/L4, A/X7, A/L41) (Supplemental Table 3). We determined the relative contribution of each allele to the overall number of peaks for each co-expressed sample by intersecting the peaks with merged peak sets for the individual variants (bedtools^63^, see Methods; Supplemental Table 6). We assigned peaks that overlapped uniquely with one of the two variants to that variant, and peaks that overlapped peaks from both variants as “shared.” Finally, we identified a subset of peaks (mean + SD = 42.27 +/- 17.1%) that appeared unique to the coexpressed samples. While the significance of this finding is unclear, motif enrichment analysis (homer^65^) of these peak subsets suggests that most are likely to be lower affinity sites that were not recovered in previous experiments.

We benchmarked our standard approach and analysis pipeline using combinations for which *in vivo* recombination data was available (A/B, A/C, C/L4, and A/N^29,50,61^). A/B co-expression resulted in H3K4me3 deposition at both A and B hotspots (Fig. 4A; Extended Data 6). Our results for A/N were similar, except that we identified both the predicted A (*p = 1e^-46^*) and N (*p = 1e^-136^*) motifs (homer^65^) when co-expressed, suggesting that, despite being a weak A allele, the N variant likely specifies a small number of distinct hotspots (Fig. 4A-B; Extended Data 6, Supplemental Data 2^59^). Co-expression of the A/C variants resulted in H3K4me3 deposition at both A and C sites (∼1:1 ratio), and enrichment of both A (*p = 1e^-119^*) and C (*p = 1e^-52^*) motifs in A/C peaks (Fig. 4A-B and Extended Data 6; Supplemental Table 6). In contrast, we observed little contribution of the L4 allele to hotspot H3K4me3 deposition in the C/L4 sample (Fig. 4A), confirming its classification as a weak C-like allele (Fig. 3E). These results recapitulate previous findings^29,50,61^, further demonstrating the utility of our assay to model *in vivo* dynamics in a cell-culture based system.

**Figure 4:**
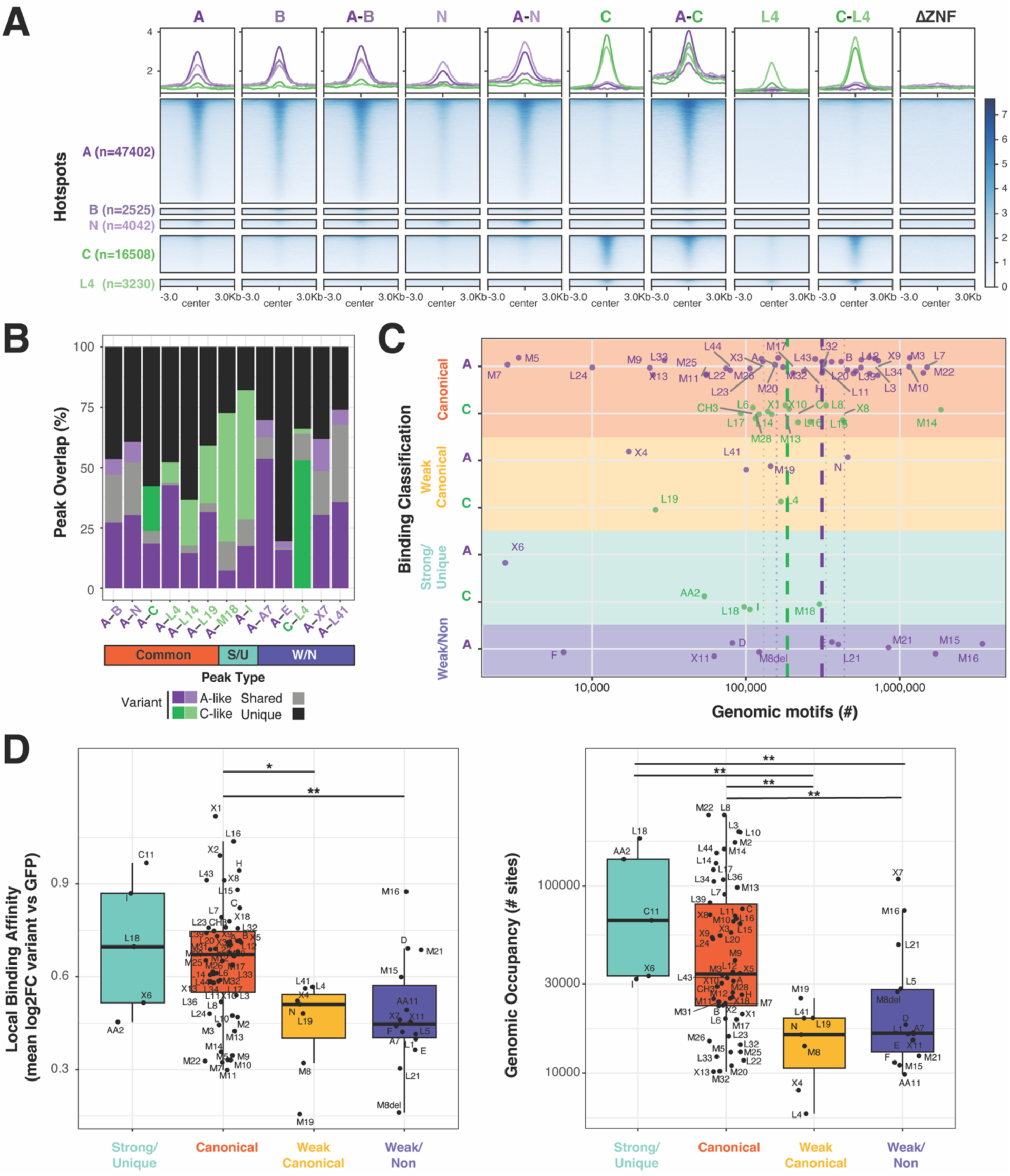
PRDM9 allelic dominance is influenced both by intrinsic molecular affinity and motif abundance. PRDM9 binding in heterozygous individuals was modeled by identifying PRDM9-dependent H3K4me3 in the presence of co-expressed PRDM9 variants. A) Heatmaps showing H3K4me3 signal at previously identified A, B, N, C, and L4 meiotic hotspots.^29,50,61^. B) Stacked bar chart showing proportion of peaks identified in co-expressed samples contributed by individual PRDM9 variants. C) Strip plot summarizing the frequency of predicted binding motifs for each variant genome-wide, categorized by allele and binding subtype. Dashed vertical lines represent the median number of predicted binding motifs genome wide for canonical A and C variants respectively, with dotted vertical lines representing the boot-strapped 95% C.I. of the median. The X-axis is log10 scaled. D-E) Boxplots summarizing local binding affinity (D) and genomic occupancy (E) (see Methods) for PRDM9 variants, grouped by binding classification. The y-axis in E is log10 scaled. Cross-group differences in local binding affinity and genomic occupancy are statistically significant (Kruskal-Wallis, df=3, *p* < 0.05). * Benjamini-Hochberg adjusted *p <* 0.05 and ** *p* < 0.01 (Dunn’s Pairwise Comparisons test).

We next considered other combinations, including high frequency (A/L4, A/L14, A/L19) and A/noncanonical allele (A/I, A/M18, A/A7, A/E, A/X7, A/L41) genotypes. Like the A/C combination, the L14 and L19 alleles contributed approximately half of the binding when co-expressed with A (∼1:1 ratio for both; Fig. 4B; Supplemental Table 6). In contrast, L4 specifies only 1/5^th^ of the A/L4 peaks, suggesting it is not dominant to the A allele (Fig. 4B). For A/noncanonical allele combinations, their binding patterns varied based on their allele binding subtype. The strong/unique binders I and M18 both specified the majority of H3K4me3 sites when co-expressed with the A allele (∼1:3 and ∼1:7.5 ratio for A/I and A/M18 respectively; Fig. 4B, Supplemental Table 6). In contrast, weak canonical and weak/nonbinder alleles A7 (∼7.5:1), E (∼5:1), X7 (∼2:1), and L41 (∼6:1) accounted for only a small percentage of peaks when co-expressed with A (Fig. 4B; Supplemental Table 6). Thus, several variants are either dominant (M18, I) or roughly equivalent to A (C, L14, L19) (Fig. 4B; Supplemental Table 6). In contrast, A was dominant to several other variants (B, N, L4, A7, E, X7, and L41) (Fig. 4B; Supplemental Table 6).

### PRDM9 allelic dominance is driven primarily by PRDM9 binding affinity

At least two explanations might account for the dominance patterns of individual PRDM9 variants: 1) some variants might bind more strongly or weakly to their target sites (ie: affinity) and/or 2) their motif may be more abundant genome-wide. To investigate these possibilities, we determined the frequency and distribution of the top scoring motif in the human genome (hg38) for all PRDM9 variants (homer^65^) (Supplemental Table 4, Supplemental Data 5^59^). Consistent with previous studies, the number of “predicted” PRDM9 binding sites was greater than the number of empirically determined sites (A *p=2.5e-9,* n=53, W = 42, df = 1; C *p = 6.68e-5,* n=21, W = 12, df=1, paired Wilcoxon tests), demonstrating that motif presence alone is insufficient to predict PRDM9 binding (Extended Data 7). For canonical PRDM9 variants, we found that the # of motifs genome-wide was similar for A and C variants (A = 95% boot-strapped CI of median [158552, 436728], C = 95% boot-strapped CI of median [130309, 332175]) (Supplemental Table 4). In contrast, most weak canonical, strong/unique, and weak/nonbinder alleles fell outside these confidence intervals (Fig. 4C; Supplemental Table 4), and genome-wide motif frequency did not always fit expectations (many motifs = strong, few motifs = weak). Motif abundance for several weak canonical (N, L4, M19) or weak/nonbinder (E, L21, M15, M21) variants is comparable to or greater than their respective canonical variants. Similarly, the motif abundance for only one strong/unique variant (M18) exceeds that of the median canonical variant. These observations suggest that differences in PRDM9 variant dominance cannot be solely explained by motif abundance.

We then quantitatively assessed the binding affinity (batch-corrected log2FC of normalized read counts [variant H3K4me3/GFP H3K4me3] within peaks) and genome occupancy (total number of peaks across replicates) for each variant (see Methods) (Fig. 4D-E, Supplemental Table 4). The binding affinity (mean ± SD) was significantly different (*p =* 2.80e^-03^, Kruskall-Wallis test, H=14.07, df=3) across binding subtypes, with canonical variants (n=54) binding more strongly (0.649 ± 0.188) than both the weak canonical (n= 7, 0.446 ± 0.152; *adj. p* = 2.98e^-02^) and weak/nonbinder variants (n=14, 0.483 ± 0.180; *adj. p* = 2.06e^-02^) (Dunn’s pairwise comparisons test, Benjamini Hochberg FDR). The strong/unique variants (n=5) also tended to bind with higher affinity (0.701 ± 0.221) than either the weak canonical or weak/nonbinder variants, though this difference was not statistically significant (*adj. p* = 6.63e^-02^; Dunn’s pairwise comparisons test, Benjamini Hochberg FDR). Genome occupancy was also significantly different (*p =* 1.16e^-03^, Kruskall-Wallis test, H=15.95, df=3) between binding subtypes, with both strong/unique (89810.5 ± 66467) and canonical variants (62724.2 ± 60249.5) binding more sites genome-wide than either the weak canonical (15521.9 ± 6751.5) or weak/nonbinder (29577.6 ± 28947.2) groups (strong vs weak canonical *adj. p* = 1.06e^-02^; strong vs weak/non *adj. p =* 2.03e^-02^; canonical vs weak canonical *adj. p* = 1.06e^-02^; canonical vs weak/non *adj. p* =2.32e^-02^; Dunn’s pairwise comparisons test, Benjamini Hochberg FDR). For both metrics, the strong/unique variants trend towards binding with higher affinity (0.701 ± 0.221 vs 0.649 ± 0.188) to more sites (89810.5 ± 66467 vs 62724.2 ± 60249.5) than the canonical variants. We observed no significant differences in binding affinity or genome occupancy between the weak canonical and weak/nonbinder groups, suggesting these variants have comparable binding affinity. These results are broadly consistent with the results of our co-expression assay and suggest that the primary factor in determining PRDM9 allelic dominance is binding affinity.

### PRDM9 variants identified in infertile individuals are functional outliers with two binding modes

Having defined the functional landscape of human PRDM9 allelic variation, we next investigated how variants identified in infertile individuals (Fig. 1C) compared to canonical ones. Of the minor variants identified in our NOA and POI cohorts, most were also identified in our fertile controls (B, C, L20, L24, M19) and/or bind similarly as other canonical variants (B, C, L20, L24, M19, M10, L11), suggesting they are unlikely to impair fertility. We therefore focused on five remaining variants (A7, L7, E, M8del, and M18), all of which are heterozygous with the A allele (Supplemental Table 2). Except for L7, all are noncanonical alleles. Two variants (L7 and M18) bind strongly to many sites, and the remainder (A7, E, and M8del) bind weakly to fewer sites (Fig. 5A). Additionally, the local binding affinity of most of the infertility-identified variants are ranked near the top (M18 = 4/79, L7 =11/79) or bottom (A7 = 66/79, E = 68/79, and M8del =78/79) of the distribution (Supplemental Table 4), suggesting they are functional outliers.

**Figure 5:**
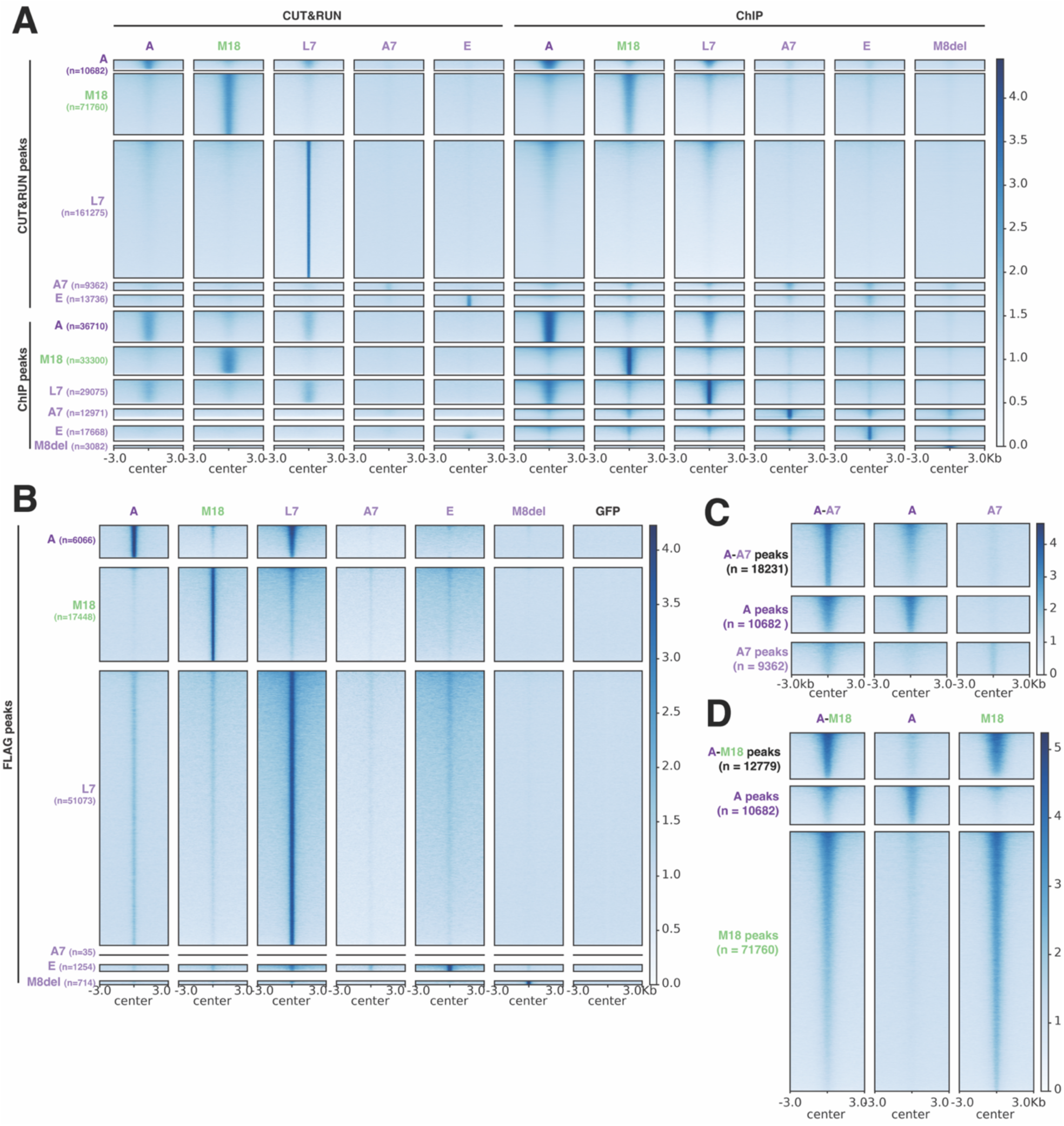
PRDM9 alleles identified in infertile individuals are functionally distinct from canonical PRDM9 alleles. A-B) Heatmap of H3K4me3 (CUT&RUN/ChIPseq; A) or FLAG signal (ChIPseq; B) over peaks called for the A variant or four patient variants. C-D) Heatmap summary of co-transfection experiments for the patient A7 (D) and M18 **variants (E). Heatmap binding signals are represented as ratios (PRDM9-variant/PRDM9-ΔZNF control for CUT&RUN, PRDM9-variant/input for ChIPseq).**

To verify that these differences in H3K4me3 deposition are due to binding properties of the ZNF array as opposed to differences in methyltransferase activity, we compared direct binding (anti-FLAG ChIPseq) of M18, L7, A7, E and M8del alleles (Fig. 5B) to several variants identified in the fertile controls (A, C, L20, L24, and M19) (Extended Data 8). In all cases, the anti-FLAG ChIP identified fewer bound regions, though the FLAG and H3K4me3 datasets were highly correlated (Pearson *r* = 0.61 – 0.88; Fig. 5C, Extended Data 4). While this discrepancy is likely influenced by antibody sensitivity, it may also reflect differences in PRDM9 activity, as deposited H3K4me3 may be more durable (and thus detectable at more sites) than direct PRDM9 binding. Consistent with this, direct binding of the weaker patient variants was minimal (A7 n=35, E n=1254, and M8del n=714 peaks respectively) while the stronger variants bound more or comparably (M18 n=17448, L7 n=51073) to alleles identified in the fertile cohort (A n=6066, C n=38085, L20 n=12355, L24 n=9052, M19 n=2598) (Extended Data 8). Collectively, these results confirm the binding classifications of M18 (strong/unique) and A7/E/M8del (weak/non) and suggest differences in PRDM9-dependent H3K4me3 deposition are driven by divergent DNA binding capabilities rather than methyltransferase activity.

Finally, to interrogate how infertility-identified variants might affect PRDM9 binding in heterozygous individuals, we replicated three patient genotypes (A/A7, A/E, and A/M18) using our co-expression assay. For both the A/A7 (Fig. 5C) and A/E (Extended Data 8) combinations, while there is some H3K4me3 deposition at the minor variant sites in the co-expressed samples, almost all co-expressed peaks (A/A7 or A/E) are driven by the major A allele. In contrast, the A/M18 peaks are almost completely determined by the M18 allele, with M18 redistributing the A allele to its own sites (Fig. 5D). These results collectively suggest two binding modes for infertility-identified variants: null (M8del, A7, E) and gain of function (M18, L7).

## Discussion

Although infertility is common^1^ and poses a steep personal and economic cost^3^, the underlying etiology is unknown in up to 30% of cases^2^. Because defects in MR contribute to infertility, proteins that regulate MR have been identified as possible infertility genes. PRDM9, encoded by one the most rapidly evolving vertebrate genes^31^, is the primary determinant of MR hotspot locations in humans^10–12^. Despite decades of research identifying novel human PRDM9 alleles^10,12,29–31,54,55^, the functional consequences of this extensive allelic variation has remained largely unknown due to difficulties sequencing *PRDM9*’s ZNF array, a coding minisatellite, and the resource-intensive nature of traditional DNA binding profiling assays (ie: ChIPseq). While long-read sequencing^29^ has substantially expanded the catalog of PRDM9 alleles, most were identified in individuals of unknown fertility status, precluding determination of whether these alleles represent normal variation or alter hotspot specification sufficiently to affect MR. Here we interrogated the functional consequences of PRDM9 variation, including its contribution to human infertility by 1) characterizing the existing functional variation, including binding affinity and bound genomic sites/motifs, for 80 human PRDM9 variants, 2) identifying PRDM9 alleles from individuals with known fertility status (fertile men, men with NOA, and women with POI) using long-read sequencing, and 3) using this data to contextualize infertility-identified variants.

Surprisingly, our data reveals a discordant relationship between human PRDM9 genetic and functional diversity. Even PRDM9 variants that differ markedly in zinc-finger number and sequence often behave indistinguishably from the canonical A or C alleles, including binding nearly identical DNA motifs and loci. Thus, rather than continuously exploring new DNA binding specificities, most segregating PRDM9 alleles preserve one of two successful hotspot-specification programs. However, variants can escape these two binding modes via two distinct trajectories: 1) by establishing new hotspot programs through novel DNA binding specificity (strong/unique binders) or 2) reducing overall hotspot number via loss or reduction of DNA binding affinity (weak/nonbinders). Historical recombination (*r*) patterns support this interpretation: sites specified via canonical alleles display high *r,* consistent with long-term usage, whereas sites specified by novel binders show intermediate *r,* suggestive of recently established hotspot programs, and weak/nonbinder variants specify sites with minimal *r* above background. This framework offers a functional perspective on the rapid evolution of PRDM9, aligning with computational models^44,45^ that predict continuous hotspot turnover due to meiotic drive, which erodes high-affinity PRDM9 binding sites and favors evolution of PRDM9 alleles with new DNA-binding specificities to restore sufficient crossover numbers to complete MR. Collectively, our results suggest that human PRDM9 evolution is highly canalized, with relatively few functional outcomes tolerated despite frequent coding sequence changes.

Although our study was neither designed nor sufficiently powered to causally link PRDM9 genotype to infertility in our NOA and POI cohorts, our results nonetheless demonstrate that some infertile individuals possess rare alleles that are either presumptive gain of function (L7, M18) or null (A7, M8del, E) in our assay. Despite their contrasting molecular properties, we predict that either class of functional outlier would reduce symmetric PRDM9 binding between homologous chromosomes when paired with the A allele, as observed in our patients (Fig. 6A-B). Symmetric PRDM9 binding, which facilitates homolog pairing and efficient double stranded DNA break repair, has emerged as a critical determinant of successful meiotic recombination^23–28,33^. We therefore propose that disruption of PRDM9 binding symmetry represents a unifying mechanistic framework through which both gain-of-function and null PRDM9 alleles may converge on a common biological consequence (ie: infertility), though determining causality will require future studies capable of directly linking PRDM9 genotype to MR hotspot locations.

**Figure 6:**
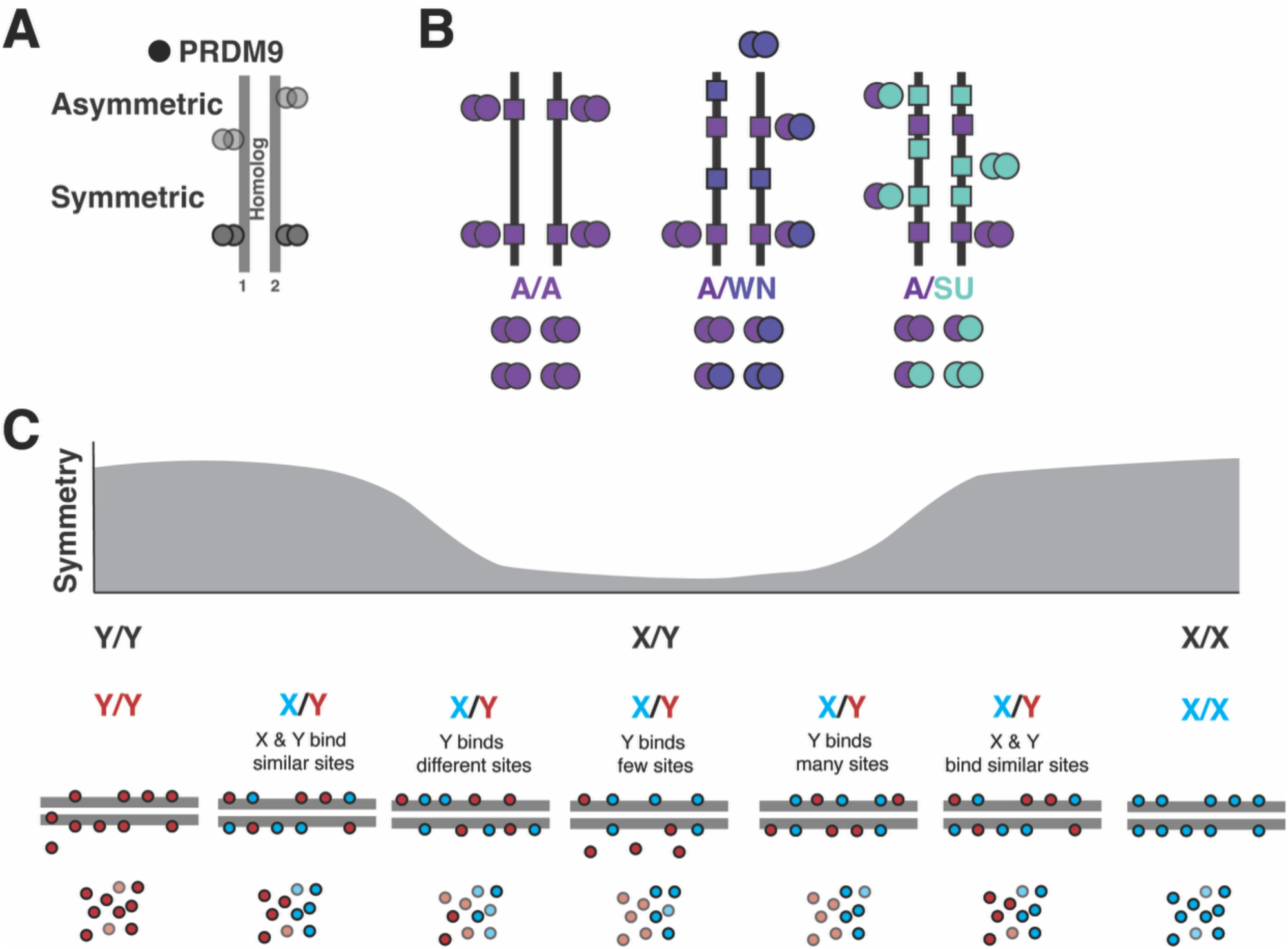
Incompatible PRDM9 variants likely increase PRDM9 binding asymmetry. A) Symmetric PRDM9 binding occurs when PRDM9 molecules (dark circles) bind to the same ∼1kb window on both parental homologs, whereas asymmetric PRDM9 binding occurs when PRDM9 molecules (faded circles) bind only one parental homolog within the same window. B) A conceptual model of how variants identified in infertile individuals might affect PRDM9 binding symmetry. In an A/A individual, symmetry is predicted to be high due to all PRDM9 molecules binding the same motif, though individual sites might vary depending on the genetic context. Individuals with an A/WN (weak/nonbinder) genotype are predicted to have reduced PRDM9 binding symmetry due to a combination of different binding preferences and a reduction in the total number of PRDM9 molecules capable of binding DNA. Individuals with an A/SU (strong/unique) genotype are predicted to have reduced symmetric PRDM9 binding due to different binding preferences and a reduction in overall A hotspots due to the SU variant diverting PRDM9-A molecules to its own binding sites. C) A general model of how different binding specificities and affinities would be predicted to reduce PRDM9 binding symmetry in a heterozygous individual. Symmetry is predicted to be high in homozygous XX or YY individuals, while X/Y heterozygous individuals may have a range of PRDM9 binding symmetry depending upon how different the binding properties are for each variant.

Our findings also help reconcile seemingly contradictory observations from mouse and human genetics. In mice, while *Prdm9* loss causes severe infertility on the C57BL/6J background^16,17,70^, fertility can be restored in certain hybrid backgrounds^70^, highlighting the importance of genetic context. In humans, while rare heterozygous *PRDM9* mutations outside the zinc-finger array, including variants within the SET domain, have been associated with NOA and POI^18–22,71^, at least one individual homozygous for an apparent *PRDM9* null allele has reproduced^72^. Collectively, this data and our own observations suggest that PRDM9 function exists along a continuum determined by the intersection of both intrinsic (DNA binding specificity and affinity; methyltransferase activity) and extrinsic (allelic dosage, genetic background, motif abundance, hotspot symmetry) factors. Thus, while we propose that the alleles we identified in our infertile cohorts might be incompatible with the A allele due to reduced symmetric binding, they might not impact fertility in all cases (Fig. 6C).

Our research also provides a practical framework for interpreting newly discovered PRDM9 variants. Due to recent advances in long-read sequencing and PRDM9’s elevated mutation rate, which is estimated to be 1-2 orders of magnitude greater than that of typical coding loci^31^, future studies will undoubtedly identify new PRDM9 alleles. Because most individual alleles are rare, establishing statistically significant associations with infertility will require exceptionally large patient and control cohorts. Functional classification, therefore, provides an orthogonal approach for variant interpretation. Our assay, which enables high-throughput functional characterization of PRDM9 activity in a tractable cell culture system that recapitulates many features of *in vivo* PRDM9 binding, can thus be used to classify novel variants into defined functional categories (canonical, weak canonical, strong/unique, and weak/non-binding). Such annotation will facilitate prioritization of candidate pathogenic alleles for further genetic and mechanistic studies.

Finally, our work highlights fundamental questions regarding PRDM9 biology that remain unresolved. Our results indicate that hotspot specification is determined by numerous factors, including ZNF sequence, genomic motif abundance, intrinsic DNA binding affinity, allelic dominance in heterozygous individuals, and overall PRDM9 expression. However, the quantitative requirements of this process are unknown, such as how many PRDM9 molecules are present within the meiotic nucleus, how many sites need to bound and modified by PRDM9 to ensure successful recombination, and whether a threshold number of PRDM9-marked hotspots is necessary for fertility. Likewise, how variation in PRDM9 occupancy influences recruitment of downstream factors such as ZCWPW1 and ZCWPW2, which recognize the dual H3K4me3/H3K36me3 mark and couple PRDM9-dependent chromatin modification to double-stranded DNA break formation and repair^73–77^, is poorly understood. Future work addressing these questions will contextualize our results and improve the ability of both theoretical and computational models to predict the functional consequences of PRDM9 variation.

Collectively, our study, which leverages human genetics and high throughput genomics to characterize the functional diversity of human PRDM9, identified an unexpectedly constrained pathway for hotspot specification, where most alleles preserve established MR hotspots, but rare variants can radically reshape or diminish them. By linking both gain-of-function and null alleles to disruptions in hotspot symmetry, we propose a unifying mechanism whereby PRDM9’s role in facilitating MR is dependent upon its intrinsic molecular properties and the broader genomic context in which it acts. This framework improves our understanding of PRDM9’s evolutionary dynamics and enables functional classification of future PRDM9 variants. Although PRDM9 is not currently part of the standard genetic evaluation of NOA, which remains largely limited to karyotype, Y-chromosome microdeletion, and CFTR analysis^78^, its established role in murine infertility and the functional classification framework presented here provides a compelling rationale for including PRDM9 in future expanded diagnostic panels, pending validation in larger, phenotypically well-characterized cohorts.

## Methods

### Obtaining and aligning human PRDM9 ZNF arrays

To obtain the protein sequences of previously described PRDM9 ZNF arrays, we surveyed existing publications^10,12,29–31,54,55^ and queried the NCBI Genbank Nucleotide Database for deposited sequences corresponding to the last exon, which encodes the ZNF array, of the *PRDM9* gene. This approach identified 85 unique PRDM9 variants (Supplementary Table 1). For each variant, we recorded the full AA sequence, the fingerprint residue AAs, and the total number of ZNFs, including the number of predicted multimerization and DNA-binding ZNF units (Supplementary Table 1).

We sought to classify the variants by both array structure and AA similarity. To do this, we first generated a protein alignment of the arrays (MUSCLEv5.^79^). Because of the high similarity of non-fingerprint residues in PRDM9’s individual ZNF units^36^, we manually curated this preliminary alignment and present it as a visual representation of fingerprint AAs. Starting with the first ZNF of the array (fingerprint WHI), we aligned each finger in order without gaps between fingers if the fingerprint residues were either the same or differed by 1 AA (ie: WVT aligned with WVT or WVS). In the case of duplicated fingers, we aligned them to the leftmost (N-terminal) end of the alignment immediately after the original non-duplicated finger, leaving a gap for other variants in the alignment that lacked the duplication. For loss of a ZNF unit, we aligned the remaining fingers to rightmost (C-terminal) end of the alignment for which the fingerprint AAs matched, leaving gaps as necessary. Using our fingerprint AA alignments, we classified the arrays based on the C-terminal region of the ZNF array. Because of the high similarity of individual variants to either the A or C variants, and to increase visual compactness, we split the alignments into A-like and C-like subalignments.

### Identification of and DNA acquisition from NOA, POI, and fertile control individuals

We obtained purified genomic DNA from de-identified women (n=12) with premature ovarian insufficiency (POI) (Veronica Gomez-Lobo, NICHD). We obtained purified DNA from de-identified men with non-obstructive azoospermia (NOA; n=47; Don Conrad, Oregon Health & Science University) and normospermic controls (FM; n=39; Kenneth Aston, University of Utah). This study was approved by the institutional review boards of the Eunice Kennedy Shriver National Institute for Child Health and Human Development (IRB#20CH-0126), the Oregon Health and Science University (IRB #0002020243) and the University of Utah (IRB #00012049). The FM control population had a mean concentration of 116 million/ml (36-272 million/ml) and all had a concentration within the normal range (> 15 million/ml, World Health Organization, 5th edition). We purchased sperm from an additional 24 normospermic men (FM; Fairfax Cryobank), and purified DNA as previously described (Lowe Lab, Stanford University; see detailed description below).

The average age of NOA and FM cohorts at sample collection were 36 years (37/47 not reported) and 31 years (32/63 not reported) respectively (Supplemental Table 2). Exact ages of POI patients at diagnosis were unknown but all presented during adolescence and were under 19 years old at diagnosis. Ancestry was either self-reported (POI and FMs) or genetically inferred from whole-exome sequencing genotypes using EthSEQ^80^, with Phase 3 population and genotype data from the 1000 Genomes Project^69^ as a reference and restricted to variants with minor allele frequency > 0.2 within the targeted exonic regions. To compare ancestry across cohorts and to account for different reporting conventions, we collapsed self-reported and inferred ancestry into the five super populations (AFR = African, AMR = American, EUR = European, EAS = East Asian, and SAS = Southeast Asian) utilized by phase 3 of the 1000 Genomes Project^69^. We report these values as well as the original study values in Supplemental Table 2. The ancestry for men with NOA is as follows: 85.1% EUR, (40/47), 4.3% AFR (2/47), 2.1% EAS (1/47), and 8.5% not reported (NR; 4/47). For our FM cohort, which includes two datasets, the ancestry was as follows: 53.9% EUR (34/63), 1.6% EAS (1/63), 1.6% SAS (1/63), 6.3% mixed ancestry (2/63 EUR/EAS, 2/63 EUR/AFR), and 36.5% (23/63) NR. For the POI cohort, ancestry was as follows: 66.7% EUR (8/12), 25% AFR (3/12), and 8.3% mixed (1/12 AMR/EUR).

### DNA purification from sperm samples

In brief, we isolated high molecular weight genomic DNA from ∼250 µl of whole sperm following a phenol-chloroform extraction protocol. Sperm was digested in 500 µl of DNA extraction buffer (10 mM Tris-HCl, pH 8.0; 0.5% SDS) supplemented with 50 µl RNase cocktail (Ambion, cat. no. AM2286) and incubated at 37°C for 1 h with gentle rocking, followed by addition of 50 µl proteinase K (20 µg/µl; New England Biolabs, cat. no. P8107S) and a second incubation at 50°C for 2 h. Digestion viscosity was monitored to avoid DNA degradation from over digestion. Samples were centrifuged (500 × g, 2 min, room temperature) to pellet undigested material, and the aqueous layer was split between two 2 ml Phase Lock Gel tubes (5PRIME, VWR cat. no. 2302830) for parallel extraction. An equal volume of phenol-chloroform-isoamyl alcohol (25:24:1; pH 8.0) was added to each sample, mixed by gentle inversion, and centrifuged (5,000 × g, 15 min, room temperature); the aqueous phase was transferred to a fresh tube, and this extraction step was repeated for a total of three rounds to ensure removal of protein contaminants. DNA was precipitated from the final aqueous phase by adding one-ninth volume of ammonium acetate and 2.5 volumes of 100% ethanol, followed by gentle inversion until DNA precipitated. The precipitated DNA was resuspended in 200 µl TE buffer (10 mM Tris-HCl, 2 mM EDTA, pH 8.0) overnight at 4°C, with extended dissolution times as needed for complete DNA resuspension. DNA concentration and purity were assessed by Qubit BR DNA assay (Invitrogen #Q33266).

### PRDM9 ZNF array amplification

To amplify the *PRDM9* ZNF array from each NOA, POI, and the initial FM sample, we performed PCR as previously described^31^ with modifications. Nested PCR was performed using the Primestar (New England Biolabs) PCR kit. For the outer PCR reaction, we used 0.25uL each of the following primers (both 100mM): PN1.0F (ATATCCAGATCCACACAGCC) and PN2.4bR (GTGTGCAAGTGTGTGGKGAC) with the following PCR conditions: initial denaturation at 98°C for 2 minutes, [denaturing phase at 98°C for 10 seconds, annealing phase at 60°C for 15 seconds, extension phase at 68°C for 2 minutes] x 5 cycles. We then used this “outer” PCR product as input for the “inner” PCR protocol (Primestar PCR kit, 0.25uL each of the following primers: PN1.1F (GGAGCTGTAGAGTGGGAA) and PN2.4aR (GTGTGTGGKGACCACATTTG). PCR conditions for the “inner” PCR were as follows: initial denaturation at 98°C for 2 minutes followed by [denaturing phase at 98°C for 10 seconds, annealing phase at 60°C for 15 seconds, extension phase at 68°C for 2 minutes] x 25 cycles. For the additional FM samples (Fairfax Cryobank), we PCR amplified the PRDM9 ZNF array using a modified protocol: 2.5 units of cloned proofreading enzyme (PFU), 250 units of Roche Taq polymerase, 10x buffer (Applied Biosystems), 2uM dNTPs, and 100 mM primers (PN1.0F (ATATCCAGATCCACACAGCC) and R1 primer (GGGATCAGTGCGGGAAAGAA)) and 10 ng of DNA. PCR conditions: initial denaturation at 94°C for 3 minutes, [denaturing phase at 94°C for 30 seconds, annealing phase at 62°C for 30 seconds, extension phase at 72°C for 1 minute] x30 cycles. PCR products were purified using the QIAquick PCR purification kit (NOA samples and controls; Qiagen #28104) or via gel extraction (POI samples and additional controls; QIAquick Gel Extraction kit #28704) from a 1.7% agarose gel.

### PRDM9 ZNF array genotyping via long-read sequencing

To determine individual diploid *PRDM9* genotypes, we first performed either PacBio (NOA and FM samples) or Oxford Nanopore (ONT) long-read sequencing (POI samples). For PacBio sequencing, we generated single-molecule, real-time (SMRT) sequencing libraries using a minimum of 150 ng of gel-purified PCR amplified PRDM9-DNA as input according to the manufacturer’s protocol (SMRTbell prep kit 3.0; PacBio PN: 102-182-700). For the POI samples, we submitted gel-purified PCR amplified *PRDM9*-DNA for targeted long-read sequencing (Plasmidsaurus, ONT).

We analyzed the resulting long-read sequencing with two approaches. First, we filtered FASTQ reads filtered to include only those 600-1600bp in length to eliminate poor quality reads and then aligned using minimap2/2.31^81^ (`-ax map-ont` for ONT data, otherwise default parameters) to a custom genome consisting of a manually curated FASTA file of 71 known published PRDM9 alleles^29,31^. SAM files were then sorted, converted to BAM files, and summary alignment statistics were generated using SAMtools^82^. An individual was considered homozygous for a given allele if most of the reads (>90%) mapped to a single allele, and heterozygous if the percentage of mapped reads was distributed equally between two alleles. Samples with <100 reads or reads mapping in unexpected ratios were excluded. In parallel, we analyzed the long-read FASTA files using the “genotype_prdm9_LR” workflow developed by Kevin Brick (https://github.com/kevbrick/genotype_prdm9_LR) to genotype the samples^29^. Assigned genotypes (Supplemental Table 2) were verified via manual inspection of read alignments in IGV. Example code for the PRDM9 genotyping analysis is provided as Supplementary Software 1^59^.

### Generating PRDM9 expression vectors

To facilitate expression of all tested PRDM9 variants, we generated a modified pcDNA3.1(-) vector (see Supplementary Data 1 for plasmid map) to express an N-terminal FLAG-tagged human PRDM9-A variant whose sequence was modified to include an NheI restriction site upstream and XhoI restriction site downstream of its ZNF array to facilitate PRDM9 variant array swapping. We then synthesized codon-optimized versions of the ZNF array from 79 additional PRDM9 variants (Supplementary Table 1) which we subcloned into the NheI/XhoI sites (Genscript). To generate the pcDNA3.1-SR-hPRDM9-ΔZNF vector, we digested the original vector with NheI (NEB #R3131)/XhoI (NEB #R0146) restriction enzymes and performed In-Fusion cloning (Takara # 638947) to insert a small donor template containing a premature stop codon in lieu of the PRDM9-A ZNF array. We verified each vector via whole plasmid sequencing (Genscript/Plasmidsaurus).

### Cell culture and transfection

HEK293T cells (ATCC # CRL-3216) were chosen for the assay based on three parameters: 1) absence of endogenous PRDM9 expression, 2) high (>90%) transfectability using lipid-based approaches, and 3) robust expression of exogenous PRDM9 (Extended Data 1). We first cultured HEK293T cells to ∼50-70% density (37°C, 5% CO_2_) in either 12-well (CUT&RUN) or 10mm^2^ (ChIPseq) formats using high-glucose DMEM (Gibco #11965092) supplemented with 10% fetal bovine serum (Cytiva #SH30071.03) and 1X Anti-Anti (Gibco #15240062). We then transfected cells with plasmids encoding GFP (pMax-GFP [Lonza] or pSB-GFP) or a human PRDM9 variant (pcDNA3.1-hPRDM9-SR) using Lipofectamine 2000 (Invitrogen #11668027). For the co-transfection experiments, we transfected two human PRDM9 variant encoding vectors at equal concentrations. Transfected cells were harvested 24 hours later and, after confirmation of PRDM9 protein expression (western blot, see below), were processed through either our CUT&RUN (∼500,000 cells/replicate) or ChIPseq (∼20-40 million cells/replicate) pipelines (see below).

### Western Blot

To verify expression of exogenous PRDM9 protein in HEK293T cells and to assess overall stability of each PRDM9 variant, we first extracted protein from ∼2 million HEK293T cells transfected with either a GFP plasmid or one of the PRDM9 variant constructs. In brief, we pelleted transfected cells and extracted protein by lysing (5 mM Tris pH 8.0, 150 mM NaCl, 1% Triton, 1 mM DTT, 50 units/uL Benzonase [Millipore #E1014-5KU], 1X EDTA-free protease inhibitor [Roche #11873580001]) on ice, with periodic vortexing, for 30 minutes. Membranes were pelleted by centrifuging at max speed at 4°C for 10 minutes, and supernatant was collected. Protein concentration and yield were quantified using the BCA Assay (Pierce #23225). We then prepared 30∝g of protein for each sample by diluting in 1X NuPAGE LDS sample buffer (Invitrogen # NP0007) supplemented with 1X NuPAGE sample reducing agent (Invitrogen #NP0009) followed by boiling at 95°C for 5 minutes. Samples were resolved on a denaturing 4-12% Bis-Tris NuPAGE mini gel (Invitrogen #NP0323) with 1X MOPS running buffer (Invitrogen #NP0001) for 45 minutes at 200V. Proteins were transferred to a 4uM nitrocellulose membrane via wet transfer using 20% methanol/1X Transfer Buffer (Invitrogen #NP00061) at 30V for 90 minutes. Transfer efficiency was assessed via brief incubation in Ponceau S stain, followed by three 5-minute washes with TBST (0.1% Tween/TBS). Membranes were then blocked in 5% milk/TBST for 2 hours, washed three times in TBST, and then probed overnight at 4°C with anti-FLAG M2-HRP monoclonal antibody (Sigma # A8592, diluted 1:5000 in TBST). Following washes with TBST, membranes were then incubated ECL substrate (Thermo Scientific #32209) and imaged using an Azure 600 Imager (Azure Biosystems).

### CUT&RUN sample preparation

We performed CUT&RUN on cells 24 hours post transfection as previously described^60^. In brief, 500,000 cells per transfection were harvested and washed three times in 800uL Wash Buffer (20 mM HEPES pH 7.5, 150 mM NaCl, 0.5 mM spermidine [Merck MilliporeSigma #S0266-5G], and EDTA-free protease inhibitor [Roche #11873580001]) by centrifugation at 600 × g for 3 min at room temperature. Cells were resuspended in 1 mL Wash Buffer and incubated with activated Concanavalin A-coated magnetic beads (Bangs Labs #BP531; 10uL per sample, washed 3X with Binding Buffer [20 mM 1M HEPES pH 7.5, 10 mM KCl, 1 mM CaCl_2_, 1 mM MnCl_2_]) for 10 min at room temperature with gentle rotation.

Bead-bound cells were collected on a magnetic stand, cleared of supernatant, and resuspended in 100 µL Antibody Buffer (Wash buffer containing 0.025% digitonin [EMD Millipore #300410] and 2 mM EDTA) containing rabbit anti-Histone H3 (tri methyl K4) antibody (Abcam #ab8580) diluted 1:100. Cell suspensions were incubated overnight at 4°C on a nutator. Samples were then washed twice with 800 µL Digi-Wash Buffer (Wash Buffer containing 0.025% digitonin [EMD Millipore #300410]) using a magnetic separator. Beads were resuspended in 50 µL Digi-Wash Buffer containing 2.5 µL Protein A/G-Micrococcal Nuclease (pA/G-MNase; EpiCypher # 15-1016) and incubated on a rotator for 1 h at 4°C. Unbound pA/G-MNase was removed by washing the beads twice with 800 µL Digi-Wash Buffer.

Bead suspensions were resuspended in 100 µL Dig-wash buffer and chilled to 0°C on ice. Targeted digestion was initiated by adding 2 µL of 100 mM CaCl2 (2 mM final concentration) followed by brief vortexing and incubation at 0°C for 30 min. Reactions were halted by mixing with 100 µL 2X STOP Buffer (340 mM NaCl, 20 mM EDTA, 4 mM EGTA, 0.05% digitonin, 100 µg/mL RNase A [Thermo Fisher #EN0531], and 50 µg/mL glycogen [MilliporeSigma #10930193001]). Samples were incubated at 37°C for 30 min to release cleaved chromatin fragments from insoluble nuclear material, and supernatant was collected.

Resulting supernatants were treated with 0.1% SDS and 50 µg Proteinase K (2.5 µL of 20 mg/mL [ThermoFisher #EO0492) for 1 h at 50°C. DNA was isolated by adding an equal volume of phenol-chloroform-isoamyl alcohol (25:24:1), mixing, and spinning through phase-lock tubes (5PRIME, VWR cat. no. 2302830) at 16,000 × g for 5 min. Samples were further extracted with an equal volume of chloroform and centrifuged at 16,000 × g for 5 min. The aqueous phase was collected, supplemented with glycogen, and precipitated with 500 µL 100% ethanol at -20°C. Precipitated DNA was pelleted at 16,000 × g for 10 min at 4°C, rinsed with 1 mL 100% ethanol, air-dried, and resuspended in 15 µL nuclease-free water. Concentration and yield of purified DNA was quantified using the Qubit dsDNA HS assay (Invitrogen #Q32851) and used as input for library generation and downstream NGS sequencing (see below).

### ChIPseq sample preparation

20-40 million cells per sample were cross-linked (final concentration 0.75% methanol free formaldehyde; ThermoScientific #28908) in 15cm dishes for 10 minutes with gentle rotation at room temperature. We quenched the cross-linking reaction with glycine (125mM final concentration) by rotating for 5 minutes at room temperature. Media was aspirated and cells were rinsed twice with 10 mL ice cold PBS, and then cells were scraped in 8 mL ice cold Farnham Lysis Buffer (5 mM PIPES pH 8.0, 85 mM KCl, 0.5% NP-40, supplemented with fresh protease inhibitor cocktail [Roche #11873580001]) and transferred to 15 mL conical tubes on ice. Cells were pelleted at 2000 rpm for 5 minutes at 4°C, and pellets were snap frozen on dry ice and stored at - 80°C until sonication.

Cross-linked cells (∼20 million/sample) were thawed on ice for 10 min and lysed in 1 mL Farnham Lysis Buffer for 10 min on ice. Cells were homogenized with 20 strokes using a type-B dounce and pelleted by centrifugation at 2,000 rpm for 5 min at 4°C. The nuclear pellet was resuspended in 500 µL Nuclei Lysis Buffer/RIPA (50 mM Tris-HCl pH 8.0, 150 mM NaCl, 2 mM EDTA, 1% NP-40, 0.5% sodium deoxycholate, 0.1% SDS, fresh protease inhibitor cocktail [Roche #11873580001]) per 10 million cells and incubated on ice for 10 min. Chromatin was sheared to a target size of 200–500 bp using a Bioruptor Plus on high setting (30 s ON / 30 s OFF for 30 cycles, resting 5 min every 10 cycles at 4°C). Lysates were cleared by centrifugation at 20,800 × g (14,000 rpm) for 15 min at 4°C, and soluble chromatin supernatant was collected and stored at - 80°C or pre-cleared with Protein A (Invitrogen #10001D) or G (Invitrogen #10003D) Dynabeads (30 µL per sample, incubated at 4°C with end-to-end rotation for 1 hour) and used directly for ChIP (see below)

Protein A or G Dynabeads (30 µL) were washed three times with 0.8 mL cold 0.5% BSA/PBS, resuspended in 0.8 mL RIPA buffer, and incubated with 5 µg of primary antibody (rabbit anti-Histone H3 (tri methyl K4) antibody [Abcam #ab8580] or mouse anti-FLAG M2 monoclonal antibody [Sigma-Aldrich #F3165]) for 2–6 h at 4°C with rotation. Antibody-coupled beads were washed three times with 0.8 mL 0.5% BSA/PBS. Pre-cleared chromatin (100 µg; ∼20 million cell equivalents in 1 mL) was added to the prepared beads and incubated overnight at 4°C with gentle rocking.

Immune complexes were washed at 4°C for 5 min per wash with 0.8 mL of the following cold buffers: once with Low Salt Wash Buffer (0.1% SDS, 1% Triton X-100, 2 mM EDTA, 20 mM Tris-HCl pH 8.0, 150 mM NaCl), twice with High Salt Wash Buffer (0.1% SDS, 1% Triton X-100, 2 mM EDTA, 20 mM Tris-HCl pH 8.0, 500 mM NaCl), twice with LiCl Wash Buffer (250 mM LiCl, 1% NP-40, 1% sodium deoxycholate, 1 mM EDTA, 10 mM Tris-HCl pH 8.0), and twice with 1X TE Buffer (pH 8.0, without rocking). Beads were then resuspended in 200 µL freshly prepared Direct Elution Buffer (10 mM Tris-HCl pH 8.0, 0.3 M NaCl, 5 mM EDTA, 0.5% SDS) containing 2 µL RNase A (10 mg/mL) and incubated overnight at 65°C with 850 rpm shaking to reverse cross-links. Supernatants were collected using a magnetic stand, incubated with 3 µL Proteinase K (Roche #03115852001; 20 mg/mL) for 1–2 h at 55°C, and purified using a QIAquick PCR Purification Kit (input samples; Qiagen #28104) or Zymo DNA Clean & Concentrator-5 kit (ChIP DNA; Zymo D4013) according to the manufacturers’ protocols. Purified DNA was eluted in 30 µL (inputs) or 15 µL (ChIP DNA) of water. Concentration and yield of purified DNA was quantified using the Qubit dsDNA HS assay (Invitrogen #Q32851) and used as input for library generation and downstream NGS sequencing (see below).

### NGS library prep and sequencing

Sequencing libraries were prepared with the Takara ThruPLEX DNA-seq Kit (Takara # R400674) and either single (#R400695/R400697) or dual index adapters (#R400665-8) (Takara) using 10 ng of purified CUT&RUN or ChIP DNA as input. We followed the standard manufacturer’s protocol with the following modifications during the Library Amplification step: 10s extensions (CUT&RUN) or 50s extensions (ChIP) both at 72°C, 12 total library amplification cycles. Following library creation, we performed either a 0.6x/1x (CUT&RUN) or a 0.65X-0.3X double-sided size selection (ChIP) using HighPrep PCR Clean Up Kit and Purification magnetic beads (MilliporeSigma #AC-6000) and quantified final yield (Qubit dsDNA HS assay; Invitrogen #Q32851) and fragment size (Agilent TapeStation High Sensitivity D1000 assay #5067). Pooled sequencing libraries were sequenced with 50x50bp PE (CUT&RUN) or 100x100bp PE (ChIP) reads (NovaSeq 6000) to a minimum of 20 million reads per sample.

### CUT&RUN and ChIPseq data analysis

Example commands for all analyses are available as Supplementary Software 1^59^.

#### Data Processing

To assess the quality of our CUT&RUN and ChIPseq datasets and to generate files for downstream analyses, we first assessed overall read quality, including individual base quality scores, proportion of duplicate reads, and presence of adapter dimers using FastQC^83^ (default parameters). If sequencing adapters were present, we trimmed them using cutadapt^84^ (-`a AGATCGGAAGAGCACACGTCTGAACTCCAGTCA -AAGATCGGAAGAGCGTCGTGTAGGGAAAGAGTGT --overlap 6 -q 20 -j 16 -- minimum-length 25`). We then aligned the reads to the hg38 genome (GCF_000001405.40_GRCh38.p14) using either bowtie2^85^ (`bowtie2 –-local –- very-sensitive-local --no-unal --no-mixed --no-discordant -- phred33 -I 10 -X 700`, parameters obtained from^60^) for CUT&RUN samples or bwa mem^86^ (default parameters) for ChIPseq samples and converted the resulting SAM files to BAM format using samtools sort (`samtools view -b -q 30` for CUT&RUN, default parameters for ChIPseq)^82^. BAM files were indexed using samtools index (default parameters)^82^.

#### Peak Calling

To identify sites where individual PRDM9 variants bind in the genome, we called peaks using MACS2^62^ with the following parameters: for CUT&RUN we used parameters outlined as previously described^60^ (`macs2 callpeak -f BAMPE -g hs –keep-dup all -p 1e-5`) and for ChIPseq we called peaks in IP over input samples (`macs2 callpeak -t IP -c input -f BAMPE -g 2913022398`). To identify PRDM9-dependent H3K4me3 peaks, we removed peaks that overlapped with H3K4me3 peaks called from batch-matched GFP-transfected samples and annotated promoters (± 1kb centered on annotated TSS, hg38_gencode_v40) (`bedtools intersect -a variant.narrowPeak -b GFP.narrowPeak -v -wa | bedtools intersect -a - -b promoters.bed -v -wa`)^63^. For variants with more than one replicate, we generated a merged master peak file and retained peaks present in two or more samples (`bedtools merge`)^63^.

#### QC metrics

After alignment and peak calling, we calculated additional quality metrics including percentage of mapped reads as well as measures of enrichment (Fractions of Reads in Peaks [FRIP], Normalized Strand Cross-correlation coefficient [NSC] and Relative Strand Cross-correlation coefficient [RSC]) and library complexity (PCR Bottleneck Coefficient 1/2 [PBC1/2] and Non-Redundant Fraction [NRF]). Percentage of mapped reads were calculated using samtools flagstat^82^ (default parameters). FRIP was calculated by counting the number of reads overlapping called peaks (MACS2^62^, see above) for each sample (`bedtools intersect -u -a BAM -b narrowPeak - ubam | samtools view -c`) and dividing by the total number of mapped reads^63,82^. We used run_spp.R to calculate NSC and RSC (`Rscript run_spp.R -c=BAM - savp -out=cross_ppq_output.txt`; phantompeakqualtools^87^), and preseq^88^ to calculate PBC1/PBC2/NRF (`preseq lc_extrap -v -o ccurve.txt BAM`).

Although variable across samples (Supplemental Table 3), the mean ± SD alignment rate (92.85 ± 4.66%) and enrichment (FRIP = 0.39 ± 0.23; NSC = 1.36 ± 0.26; RSC = 2.60 ± 1.42) meets ENCODE standards for ChIPseq experiments (FRIP > 0.01, NSC > 1.05, RSC > 0.8)^87^. To confirm the quality of the H3K4me3 samples, we also assessed binding enrichment at annotated gene promoters (± 1kb centered on annotated TSS, hg38_gencode_v40) by shuffling peak locations for each variant followed by promoter intersection to derive an empirical *p* value (regioneR, permutation test, 10000 shuffles)^64^. Enrichment was considered significant at *p < 0.0001,* and both *p* value and Z score are reported in Supplemental Table 3.

For replicated variants, we determined replicate correlation (Pearson *r*) by taking the master merged peak files and calculating the read coverage for each sample within the peak set, excluding blacklisted regions from^89,90^, using deepTools (multiBamSummary BED-file --BED merged_peak_file.bed --bamfiles - o mbsummary.npz -bl blacklist.bed; plotCorrelation -in mbsummary.npz --corMethod pearson --skipZeros --whatToPlot scatterplot --plotNumbers --removeOutliers)^91^. We also performed correlation analyses between CUT&RUN and ChIPseq read coverage for variants tested using both methods and between matched H3K4me3 and FLAG ChIP datasets using the same parameters.

#### Generating binding signal tracks

We generated bigwig tracks for each sample by normalizing to 1X genome coverage using deepTools^91^ (`bamCoverage --bam variant.BAM -o variant_SeqDepthNorm.bw --binsize 10 -- normalizeUsing RPGC --effectiveGenomeSize 2913022398 -- ignoreForNormalization chrX chrM --extendReads 200`). We also generated ΔZNF (CUT&RUN) or input (ChIPseq) normalized bigwigs (`bigwigCompare -b1 variant_seqDepthNorm.bw -b2 Ctrl_SeqDepthNorm.bw -o variant_ratio.bw --skipZeroOverZero --binSize 10 --operation ratio`), where we represent the signal from each variant as a ratio over the appropriate control sample. For replicated variants, we generated merged bigwig tracks by first merging individual BAM files (`samtools merge`)^82^ and then generated sequence depth and ΔZNF/ratio input normalized bigwig files from the merged BAMs as described above.

#### Motif calling

We identified sequence motifs enriched (binomial *p* < *1e-20*) within our individual and replicated (where possible) GFP-filtered peak sets using homer^65^ (`findMotifsGenome.pl filtered_peaks.narrowPeak hg38 -size given - mask -len 6,8,10,12,14,16,18,20`). Because we are primarily using PRDM9-dependent H3K4me3 as a proxy for PRDM9 binding and H3K4me3 located on adjacent histones is often represented as a single peak in our dataset, we opted to use -size given as opposed to focusing the motif search on the peak summit, as is traditionally done for transcription factors. Motif lengths were chosen based on biological assumptions (1 ZNF binds 3 nucleotides^43^) and to allow for binding motifs from a range of ZNF array lengths. We reported the top scoring motif in all cases except where the top scoring motif was identified in only a small fraction of peaks (< 1%) and a similarly high scoring motif with a higher peak fraction was present in the top 5 motifs. We considered a variant to have no good motif (NGM) if there were either no significantly enriched motifs or the only identified motifs were also enriched in our negative control (ΔZNF) peaks. Complete motif data for all tested variants is provided as Supplementary Data 2.

#### Assay benchmarking

To assess the ability of our CUT&RUN and ChIPseq assays to model *in vivo* PRDM9 binding, we plotted mean H3K4me3 (CUT&RUN/ChIPseq) or FLAG (ChIPseq) signal (bigwigs) over previously published PRDM9 A, B, and C hotspots^29,50,61^ or our replicated peaks called with MACS2^62^ (see above) using deeptools^91^ (`computeMatrix reference-point --referencePoint center - b 3000 -a 3000 --skipZeros --missingDataAsZero; plotHeatmap – colorMap Blues –heatmapHeight 10`). As an orthogonal approach, we tested whether our replicated peaks for each variant (A, B, or C) significantly overlapped the same known hotspot regions using the permutation strategy described for promoter regions above (regioneR^64^, n=10000 shuffles). Finally, we queried our replicated peak files for the A, B, and C variants for their cognate motif ^11,33,48^ using our motif analysis pipeline (see above).

### ChromHMM analysis of CUT&RUN data

To systematically interrogate differences in PRDM9 variant binding, we used ChromHMM^67^ to model PRDM9 binding genome-wide. We input CUT&RUN data (BAM files) for all variants as well as matched GFP controls as data in single cell format (`java -jar ChromHMM.jar BinarizeBam hg38.txt /chromHMM_BAMs table_file.txt /binarizedBAMs`). We also incorporated custom annotation files (known PRDM9A/B/C hotspots^29,50,61^, linkage determined PRDM9-A hotspots^92^, and previously published ChIPseq-determined binding sites for the PRDM9A/B/C alelles^48,49^) in conjunction with package provided annotation files (CpG islands, Exons, Genes, Transcription End Sites, Transcription Start Sites, Transcription Start Sites + 2kb flanking region, and Repeats) to facilitate state annotation. We initially used the data to train models (`java -jar ChromHMM.jar LearnModel /binarizedBAMs /chromHMM_results #states hg38`) of various states (12, 15, 18, 21, 24, 27, and 30) and annotated the individual states for each model using our custom and pre-packaged annotation files. We ultimately chose to use a 24-state model because it recovered expected genomic regions (promoters, A hotspots, C hotspots, and non-H3K4me3 methylated regions) while minimizing the number of non-H3K4me3 states. We chose to use only CUT&RUN data for our model to ensure maximum comparability between samples. All ChromHMM output files and the table file used to specify the design are supplied as Supplementary Data 3.

### Identifying historical recombination at PRDM9 binding sites

We utilized previously published^68,93^ estimates of fine-scale historical recombination data for 26 populations, representing 5 super-populations (Africa, Southeast Asians, East Asians, Europeans, and Americans), from the 1000 Genomes Project^69^. Per-generation recombination rate (*r*) bedGraph signal tracks for all 26 populations were downloaded from^93^. Tracks for each chromosome were concatenated and converted to bigwig format (`cat *.bed | sort -k 1,1, -k2,2n - > recombination_map_hg38.sorted.bed; bedGraphToBigWig recombination_map_hg38.sorted.bed hg38.chrom.sizes.sorted recombination_map_hg38.bw`). We then used deepTools^91^ to calculate and plot (computeMatrix reference-point --referencePoint center -R ./*peak_files.bed -S ./*recombination_map_hg38.bw -b 3000 -a 3000 --skipZeros -o recomb_matrix.gz –missingDataAsZero; plotProfile -m recomb_matrix.gz -out recomb_peaks.pdf --perGroup --numPlotsPerRow 5 --legendLocation upper-right --plotHeight 15 --plotWidth 10) mean *r* at PRDM9 bound sites (GFP filtered MACS2 peaks called from H3K4me3 CUT&RUN/ChIP data, see above) for all populations for each variant. We considered there to be evidence of historical recombination at a PRDM9 variant’s binding peaks in a population if the mean *r* exceeded that of the PRDM9-ΔZNF control (*r = 2e-9*), and in a super population if at least one subpopulation met this criterion. For replicated variants, we used the replicated peak sets. Mean *r* values underlying Extended Data 5 are provided as Supplementary Data 4.

### CUT&RUN co-transfection analyses

For the co-transfection analyses, we performed all the same data processing, QC, and downstream analyses as described above for single variants. We determined the fraction of co-transfection peaks that were specified by individual variants by overlapping the peaks (`bedtools intersect`)^63^ with an inclusive (merged) peak list for the individual variants (ie: for A/C we used both A and C peak datasets). This was chosen to increase detection sensitivity for rare and low affinity peaks that might not appear in every experiment. We then intersected co-transfected peaks that overlapped with one variant of the combination with peaks from the other variant to identify shared peaks (ie: A/C peaks that intersect with both A & C are considered shared). We determined the number of unique co-transfection peaks by subtracting the peaks contributed by individual variants as well as the shared peaks from the total peaks. We then called motifs on all peaks and on unique peaks to determine if we recovered the expected motifs and to identify other possible motifs that might explain binding in the co-transfected samples. Finally, we plotted the mean H3K4me3 signal from the individual variants (merged bigwig tracks, see above) and from the co-transfection experiment over both co-transfected peaks and replicated peaks for each individual variant using deepTools^91^ as described above.

### Identifying genome-wide PRDM9 binding motif locations

To determine the maximal number of sites a PRDM9 variant might bind to genome-wide, we scanned the genome using our empirically determined motif (homer^65^ `scanMotifGenomeWide.pl motif.motif hg38 -bed motif.sites.hg38.bed`) for each variant. These sites are provided as Supplementary Data 5. We then compared the number of putative sites to the number of observed sites using a paired Wilcoxon test (considered significant if *p < 0.05*), and quantified the median number of motif sites and the 95% boot-strapped confidence interval of the median for A and C canonical variants.

### PRDM9 binding affinity and genomic occupancy analyses

To quantitatively compare the molecular properties of PRDM9 variants across all batches, we developed an analytical pipeline to calculate the local binding affinity and genomic occupancy for each variant. During development and documentation of the computational analysis pipeline, large language models (Google Gemini for Government) were used to assist with Python code debugging, memory optimization, and resolving runtime errors. All generated code logic, linear regression models for batch-effect correction, statistical post-hoc calculations, and written text were independently audited, executed and validated by the authors. The author(s) take full responsibility for the integrity, accuracy, and scientific validity of the pipeline and manuscript content. The methodology used by the pipeline is described below, and code to run the pipeline is provided as Supplementary Software 2^59^.

#### Pipeline initialization and data curation

The analytical pipeline was executed using a custom Python-based framework designed for concurrent genomic interval processing and batch-corrected signal quantification. The experimental design, which includes primary metadata (‘metadata.csv’) mapping PRDM9 variant IDs to their respective paired GFP controls and experimental batches and categorical variant annotations (‘user_features.csv’) such as PRDM9 allele type (A or C) and binding subtype (‘canonical’, ‘weak canonical’, ‘strong/unique’, ‘weak/nonbinder’) were supplied as flat text files. Categorical variant annotations were used to build an integrated feature matrix. To ensure reproducibility, all analyses used a global random seed = 42.

#### Consensus peak integration and binary masking

To establish a uniform coordinate space across all independent experiments, individual peak files (in “.bed” or “.narrowPeak” formats) were pooled. The `PyRanges`^94^ library was utilized to concatenate and merge overlapping intervals, incorporating an optional genomic extension parameter (slack) to generate a master consensus peak matrix.

Following consensus generation, structural binary masks were constructed for each individual sample. Using high-speed interval tree cross-referencing, individual variant peak calls were mapped against the master consensus matrix. This resulted in a contiguous sample-by-peak binary matrix, where an integer value of 1 indicated a sample-specific peak intersecting the master region, and 0 indicated the absence of a peak.

#### Parallel read quantification and normalization

Read quantification was executed via an asynchronous multiprocessing pool, dynamically scaling to either the allocated SLURM task cores (“$SLURM_CPUS_PER_TASK”) or the maximally available system CPUs. Raw BAM files were parsed concurrently using `pysam` (https://github.com/pysam-developers/pysam) to count mapped reads within the boundaries of the master consensus peaks.

Raw read counts were first normalized to library depth, yielding Reads Per Million (RPM). To quantify binding intensity relative to internal batch controls (GFP), a pseudo-counted Log2 Fold Change (L2FC) was computed for each peak region using the following formula:

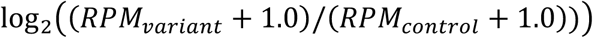

#### Network-based batch correction and affinity scoring

To mitigate technical variability across multi-batch experiments, a global regression network was deployed. The system leveraged overlapping "anchor" variants—replicated biological samples present in multiple independent batches. A multiple linear regression model (`sklearn.linear_model.LinearRegression`)^95^ was fitted using dummy-encoded variant and batch variables against the mean L2FC values of the samples. The resulting batch-effect coefficients were subtracted from the uncorrected L2FC matrix to yield a globally calibrated signal matrix. Two primary metrics were then extracted for each variant:

• *Genomic Occupancy:* the total sum of PRDM9 bound sites derived directly from the binary structure mask for each variant

• *Local binding affinity:* The median calibrated L2FC score, strictly filtered to only include regions where the PRDM9 variant’s binary mask = 1

#### Statistical evaluation and post-hoc analysis

Differences in local binding affinity and genomic occupancy across categorical feature groups were assessed using non-parametric statistical methods. A Kruskal-Wallis H-test was deployed via `scipy.stats`^96^ to evaluate global significance among groups containing at least two variants.

If the global test reached statistical significance (*p* < 0.05), post-hoc pairwise Dunn’s tests were conducted using the `scikit_posthocs` module^97^. To rigorously control for Type I errors across multiple comparisons, *p*-values were evaluated corrected using the Benjamini-Hochberg False Discovery Rate (BH-FDR) and considered statistically significant if *p* < 0.05.

#### Data visualization

Distributions of affinity and occupancy were visualized utilizing `matplotlib`^98^ and `seaborn`^99^ as side-by-side boxplots overlayed with jittered individual data points (`stripplots`).

### Statistics, visualization, and plotting

Except where outlined above, all statistical analyses and plotting were performed in R using base or published packages (ggplot2, ggpubr, ggrepel, tidyverse, dplyr).

## Acknowledgements

This work was supported by the Eunice Kennedy Shriver National Institute of Child Health and Human Development Intramural Program to T.S.M (1ZIAHD008933-13) and V.G-L (1ZIAHD009006-04) and by the Eunice Kennedy Shriver National Institute of Child Health and Human Development to D.F.C. and K.I.A (R01HD078641) and to D.F.C (P50HD096723). R.L.C. is supported by the Eunice Kennedy Shriver National Institute of Child Health and Human Development (1K99HD115747).

## Data Availability

Processed sequencing data is available at GSE345507 (ChIP) and GSE345508 (CUT&RUN). Raw sequencing data is available in the SRA. Plasmid maps, motif enrichment data, chromHMM analysis output, historical recombination data, and predicted PRDM9 motif locations are available as Supplemental Data^59^. A7 and M8del ZNF array protein sequences are deposited in Genbank (PZ704487-88). All other data is reported herein. PRDM9-expression plasmids are available upon request.

## Code Availability

Example code for all analyses and the full affinity analysis pipeline is available as Supplemental Software^59^. Exact commands used for all samples are available upon request.

---

**Extended Data 1:**
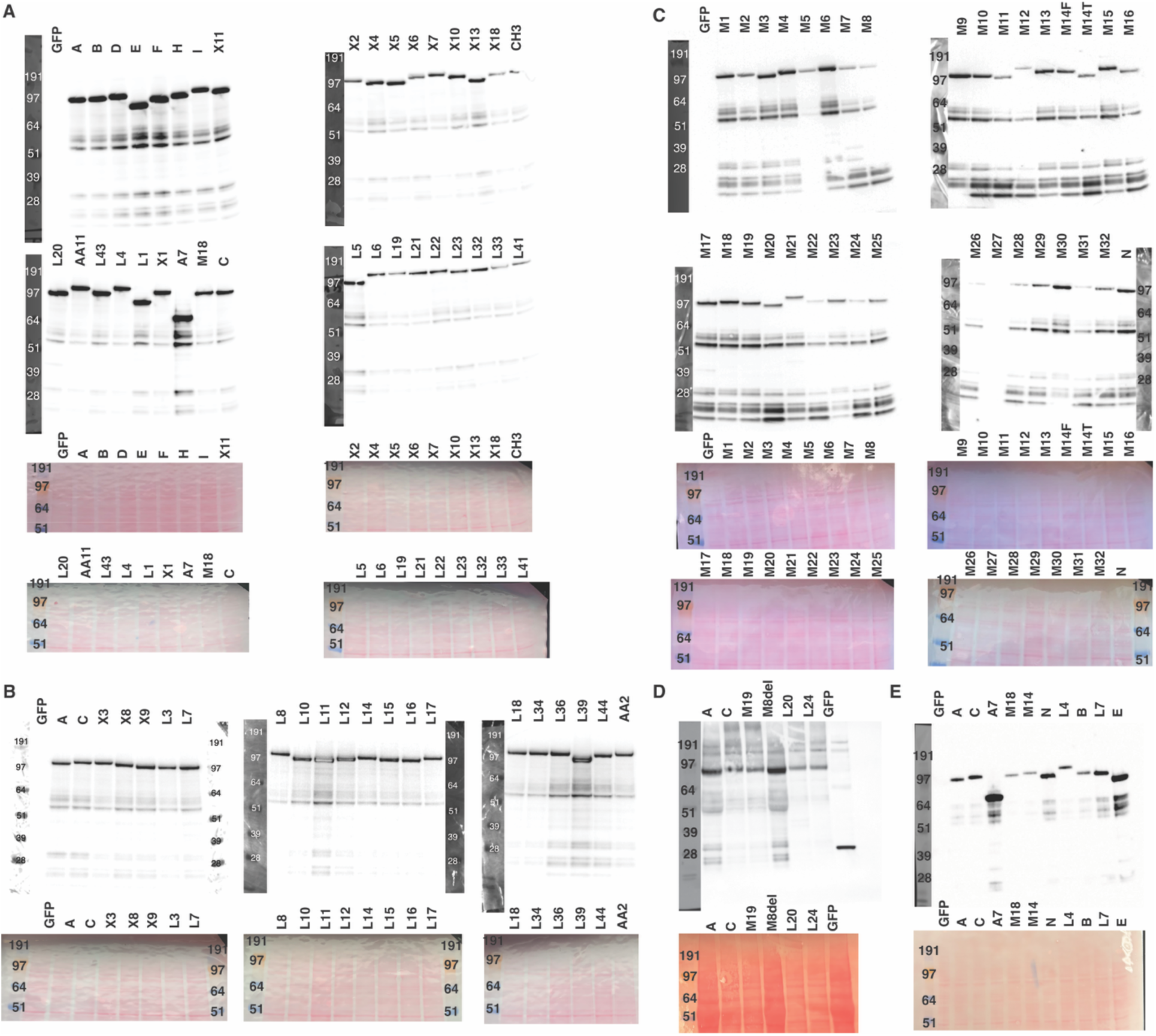
Human PRDM9 variants are expressed at similar levels in 293T cells. Protein was extracted from 293T cells transfected with vectors encoding one of the tested hPRDM9-FLAG tagged variants. Transfer efficiency/protein loading was assessed via Ponceau S stain (bottom) and then probed with anti-FLAG primary antibodies (top). A-E represent different WB experiments with associated GFP-transfected controls. A-C include variants tested via CUT&RUN and D-E include variants tested via ChIPseq. PRDM9-A migrates at ∼97 kDA.

**Extended Data 2:**
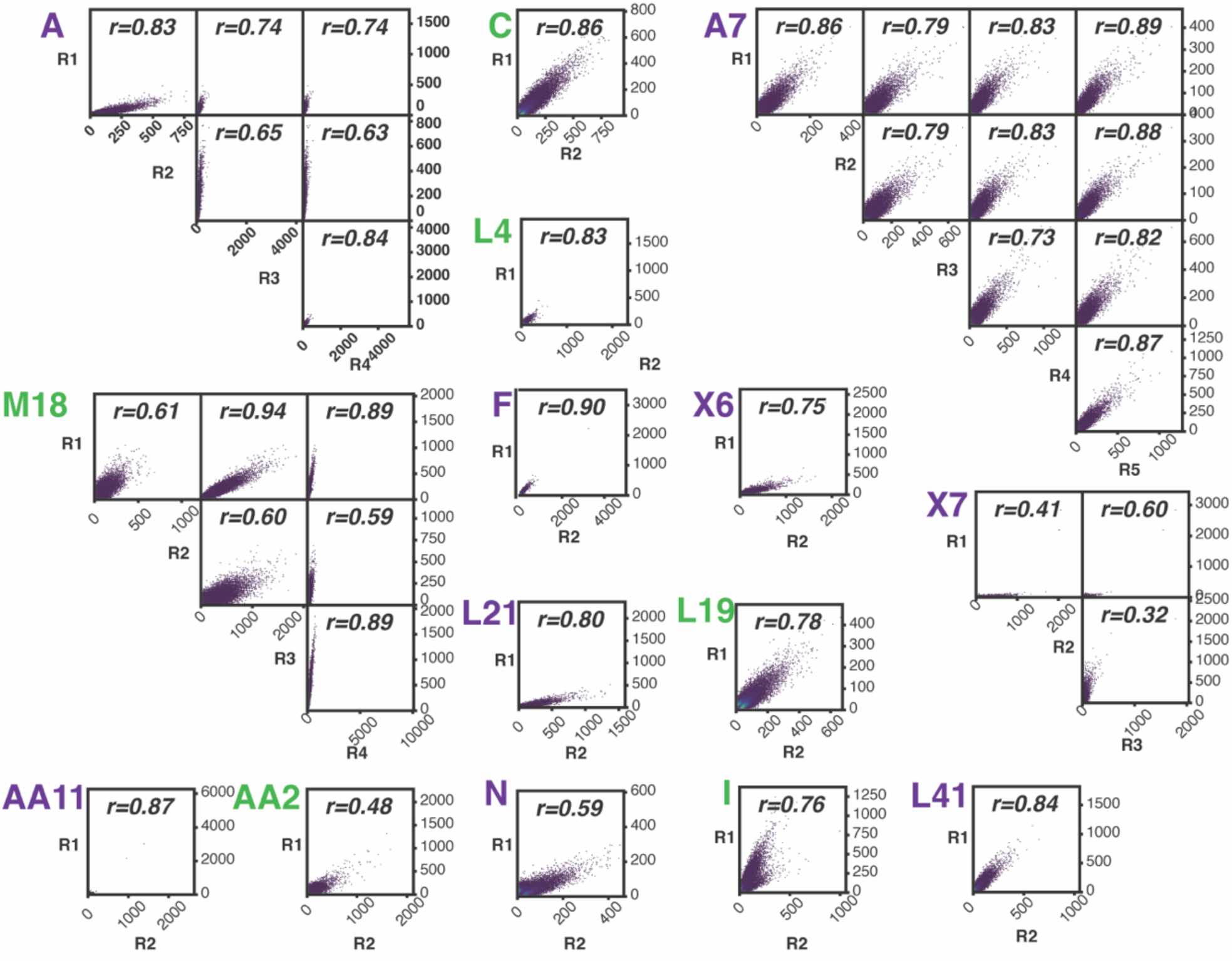
CUT&RUN PRDM9 replicate correlation. For all replicated PRDM9 variants (n > 1; CUT&RUN), we created a merged peak set and calculated the correlation (Pearson *r*) between read counts for individual replicates (R) within the merged peak set (see Methods).

**Extended Data 3:**
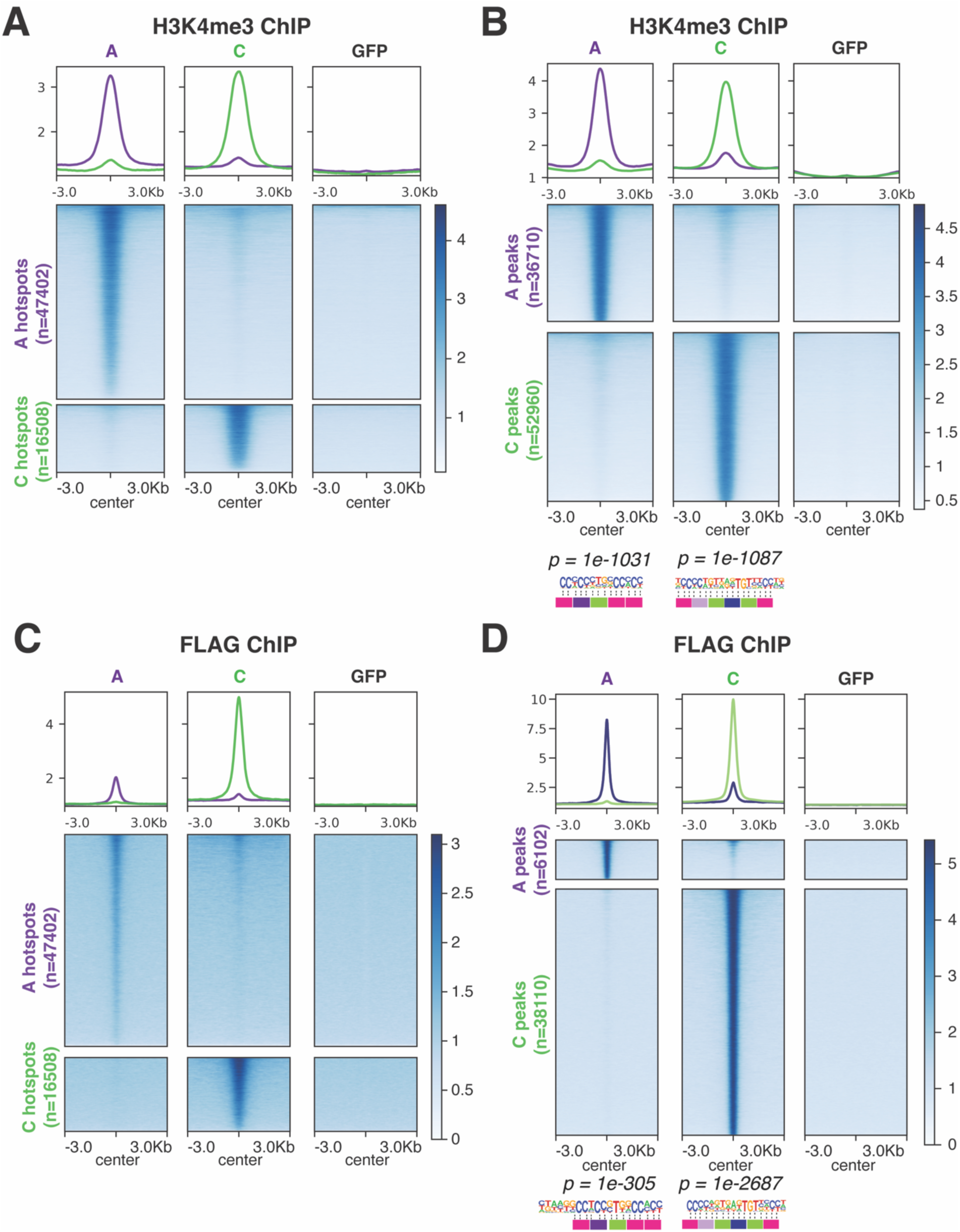
anti-H3K4me3 and anti-FLAG ChIP-sequencing assays recapitulate features of *in vivo* PRDM9 binding. A) Heatmaps plotting mean H3K4me3 signal at previously identified A and C meiotic hotspots^29,50,61^ in the presence of PRDM9 A or C. Results are contrasted with GFP as a negative control. B) Heatmaps plotting mean H3K4me3 signal at A and C H3K4me3-ChIP peaks, with logos showing the most enriched motif sequence within each variant’s respective peaks shown below. C-D) Heatmaps plotting mean FLAG signal at previously identified hotspots^29,50,61^ (C) or FLAG-ChIP peaks (D) Sequence logos in (D) show the most enriched motif sequence within each variant’s peaks. *P* values are derived from the binomial distribution.

**Extended Data 4:**
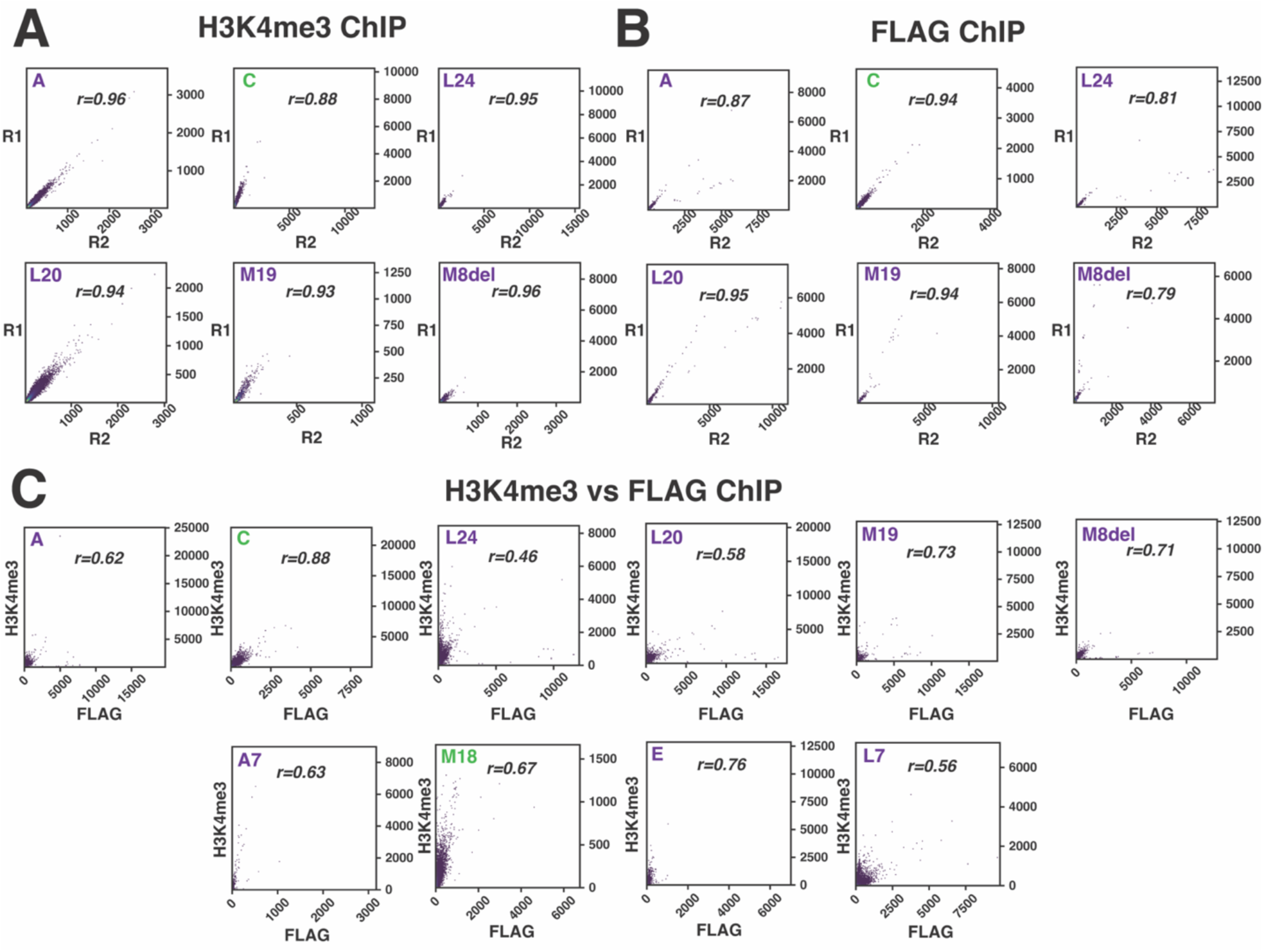
PRDM9-dependent H3K4me3 and direct PRDM9 binding (anti-FLAG) ChIP sequencing are correlated. For all PRDM9 variants tested via ChIP sequencing, we created a merged peak set and calculated the correlation (Pearson *r*) between read counts for individual replicates within the merged peak set (see Methods). A-B) Pearson *r* for all anti-H3K4me3 (A) or anti-FLAG (B) ChIP-seq replicates. C) For each variant, the Pearson *r* of read counts from H3K4me3 and FLAG samples within a peak set that combines all H3K4me3 and FLAG peaks.

**Extended Data 5:**
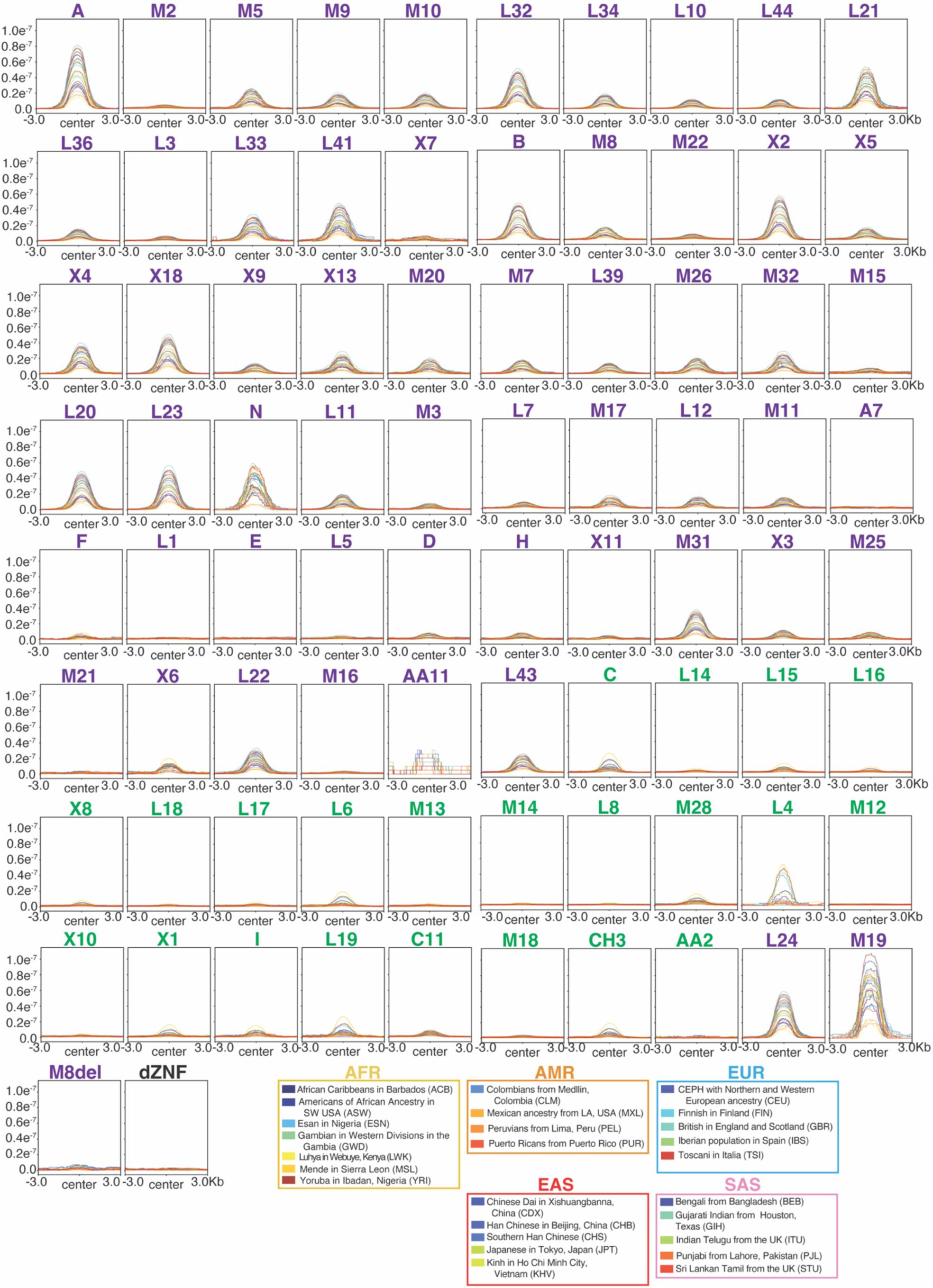
Historical recombination rate at PRDM9 binding sites. Profile plots summarizing the mean per-generation recombination rate (*r*)^68^ for 26 human populations (5 super populations: AFR = African, AMR = American, EUR = European, EAS = East Asian, and SAS = Southeast Asian) from phase 3 of the 1000 Genomes Project^69^ over 6 kb windows centered on peaks bound by each PRDM9 variant. Variant names are color coded according to allele type (A-like = purple, C-like = green).

**Extended Data 6:**
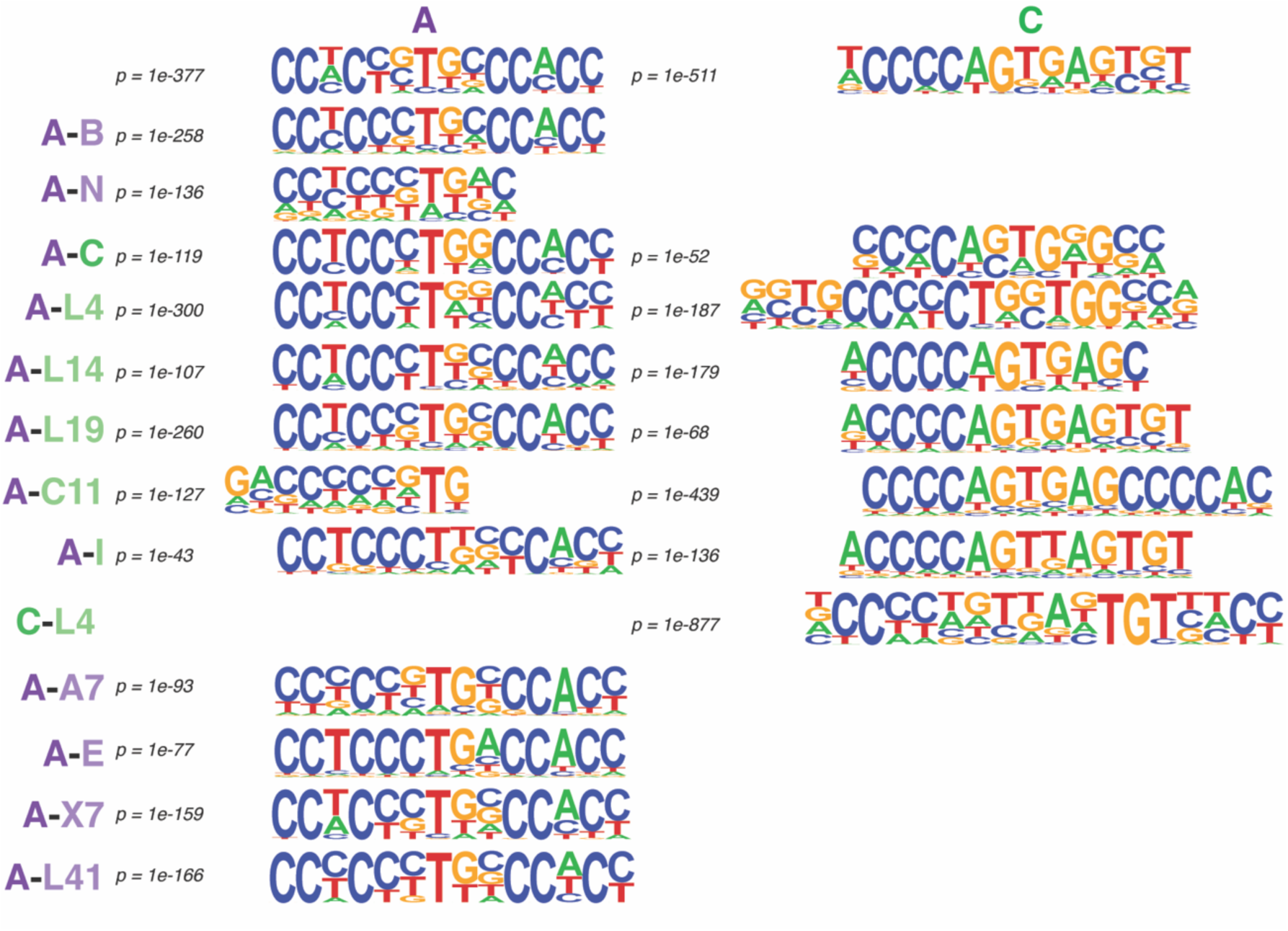
PRDM9 variant co-transfection recovers known PRDM9 motifs. PRDM9 variants shown above were co-transfected in 293T cells. Sequence logos represent the top scoring motifs within H3K4me3 peaks called in the co-transfection experiments and match motifs previously identified in this study (Fig. 3) or elsewhere^11,33,48,50^. In all cases the shown motifs were within the top 5 scoring motifs. *P* values are derived from the binomial distribution.

**Extended Data 7:**
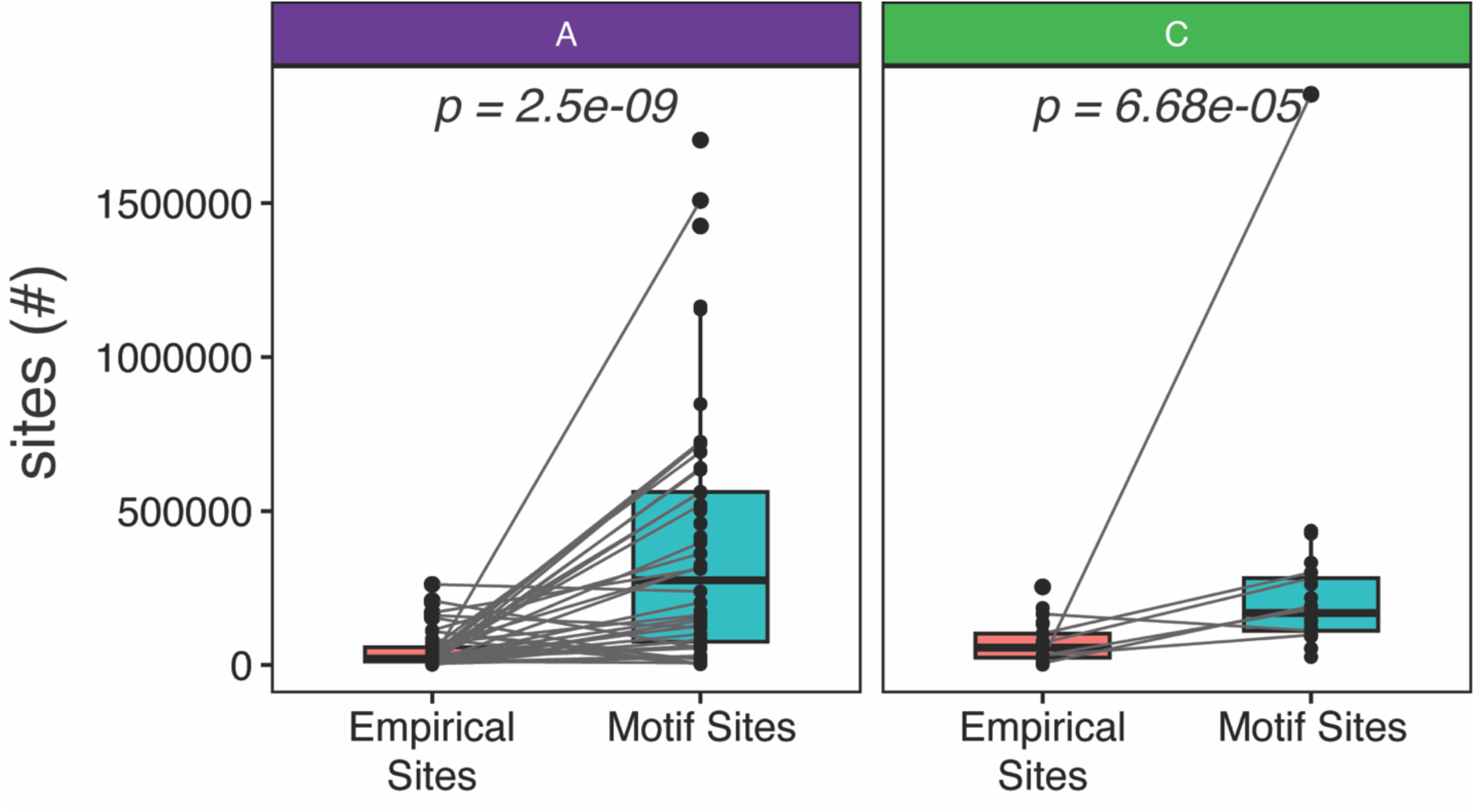
PRDM9 motifs are not sufficient for PRDM9 binding. Boxplot summarizing the distribution of empirical (ie: CUT&RUN/ChIP determined) or predicted motifs (motif sites) for each variant, faceted by type (A or C). *P* values are derived from a paired Wilcoxon test, A (n=53, W = 42, df =1), C (n=21, W = 12, df = 1).

**Extended Data 8:**
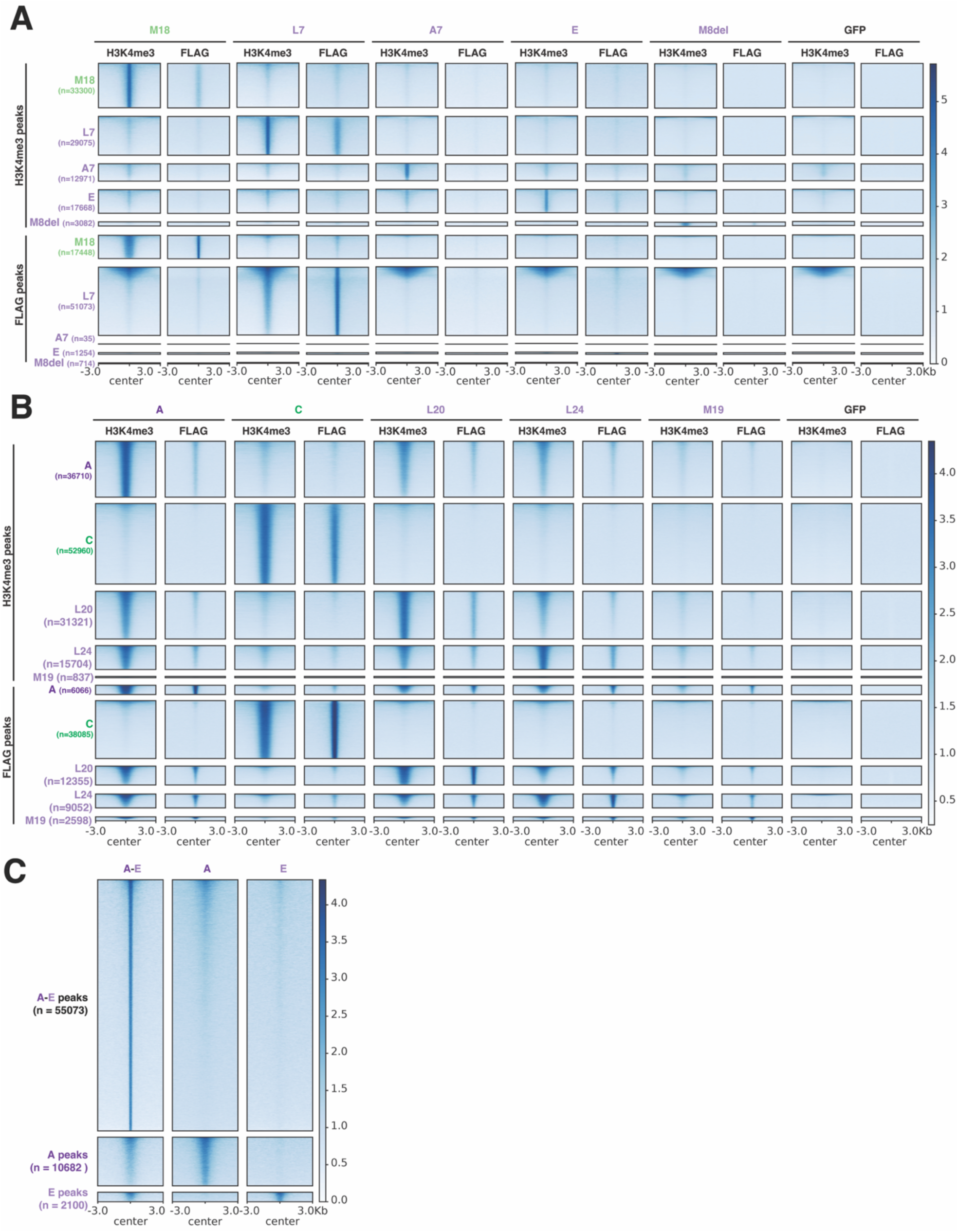
PRDM9 alleles identified in infertile individuals are functionally different than canonical PRDM9 alleles. A-B) Heatmaps of mean H3K4me3 or FLAG signal (ChIPseq) over PRDM9 variant peaks for variants identified in infertile (A) or fertile (B) cohorts. C) Heatmap of mean H3K4me3 (CUT&RUN) over peaks identified in cells expressing both PRDM9-A and PRDM9-E compared to signal at replicated PRDM9-A and PRDM9-E peaks (derived from individual expression experiments).

## References

1. World Health, O. Infertility Prevalence Estimates, 1990–2021. (World Health Organization, Geneva, 2023).

2. Chandra, A., Copen, C.E. & Stephen, E. H. Infertility and impaired fecundity in the United States, 1982-2010: data from the National Survey of Family Growth. Natl Health Stat Report, 1–18, 1 p following 19 (2013).

3. Njagi, P. et al. Financial costs of assisted reproductive technology for patients in low- and middle-income countries: a systematic review. Human Reproduction Open 2023(2023).

4. PiccheXa, L. et al. Maternal age and genome-wide failure of meiotic recombination are associated with triploid conceptions in humans. The American Journal of Human Genetics 112, 2665–2678 (2025).

5. Marston, A.L. & Amon, A. Meiosis: cell-cycle controls shuffle and deal. Nature Reviews Molecular Cell Biology 5, 983–997 (2004).

6. Rodrigo, L. et al. Sperm chromosomal abnormalities and their contribution to human embryo aneuploidy. Biology of Reproduction 101, 1091–1101 (2019).

7. Chatziparasidou, A., Christoforidis, N., Samolada, G. & Nijs, M. Sperm aneuploidy in infertile male patients: a systematic review of the literature. Andrologia 47, 847–860 (2015).

8. Arter, M. & Keeney, S. Divergence and conservation of the meiotic recombination machinery. Nature Reviews Genetics 25, 309–325 (2024).

9. Johnston, S.E. Understanding the Genetic Basis of Variation in Meiotic Recombination: Past, Present, and Future. Molecular Biology and Evolution 41(2024).

10. Baudat, F. et al. PRDM9 Is a Major Determinant of Meiotic Recombination Hotspots in Humans and Mice. Science 327, 836–840 (2010).

11. Myers, S. et al. Drive Against Hotspot Motifs in Primates Implicates the *PRDM9* Gene in Meiotic Recombination. Science 327, 876–879 (2010).

12. Parvanov, E.D., Petkov, P.M. & Paigen, K. PRDM9 Controls Activation of Mammalian Recombination Hotspots. Science 327, 835 (2010).

13. Grey, C., Baudat, F. & de Massy, B. PRDM9, a driver of the genetic map. PLOS Genetics 14, e1007479 (2018).

14. Eram, M.S. et al. A Potent, Selective, and Cell-Active Inhibitor of Human Type I Protein Arginine Methyltransferases. ACS Chemical Biology 11, 772–781 (2015).

15. Powers, N.R. et al. The Meiotic Recombination Activator PRDM9 Trimethylates Both H3K36 and H3K4 at Recombination Hotspots In Vivo. PLOS Genetics 12, e1006146 (2016).

16. Hayashi, K., Yoshida, K. & Matsui, Y. A histone H3 methyltransferase controls epigenetic events required for meiotic prophase. Nature 438, 374–378 (2005).

17. Hayashi, K. & Matsui, Y. Meisetz, A Novel Histone Tri-Methyltransferase, Regulates Meiosis-Specific Epigenesis. Cell Cycle 5, 615–620 (2006).

18. Irie, S. et al. Single-Nucleotide Polymorphisms of the PRDM9 (MEISETZ) Gene in Patients With Nonobstructive Azoospermia. Journal of Andrology 30, 426–431 (2009).

19. Miyamoto, T. et al. Two single nucleotide polymorphisms in PRDM9 (MEISETZ) gene may be a genetic risk factor for Japanese patients with azoospermia by meiotic arrest. Journal of Assisted Reproduction and Genetics 25, 553–557 (2008).

20. Soleymani Moud, S., Kamali Seraji, K., Ramezani, M. & Piravar, Z. Association of Single Nucleotide Polymorphisms in the PYGO2 and PRDM9 Genes with Idiopathic Azoospermia in Iranian Infertile Male Patients. Iranian Journal of Medical Sciences 48, 77–84 (2023).

21. Wang, Y. et al. Pathogenic variants of meiotic double strand break (DSB) formation genes PRDM9 and ANKRD31 in premature ovarian insufficiency. Genetics in Medicine 23, 2309–2315 (2021).

22. Heddar, A. et al. Genetic landscape of a large cohort of Primary Ovarian Insufficiency: New genes and pathways and implications for personalized medicine. eBioMedicine 84, 104246 (2022).

23. Smagulova, F., Brick, K., Pu, Y., Camerini-Otero, R.D. & Petukhova, G.V. The evolutionary turnover of recombination hot spots contributes to speciation in mice. Genes & Development 30, 266–280 (2016).

24. Li, R. et al. A high-resolution map of non-crossover events reveals impacts of genetic diversity on mammalian meiotic recombination. Nature Communications 10, 3900 (2019).

25. Davies, B. et al. Re-engineering the zinc fingers of PRDM9 reverses hybrid sterility in mice. Nature 530, 171–176 (2016).

26. Gregorova, S. et al. Modulation of Prdm9-controlled meiotic chromosome asynapsis overrides hybrid sterility in mice. eLife 7, e34282 (2018).

27. Hinch, A.G. et al. Factors influencing meiotic recombination revealed by whole-genome sequencing of single sperm. Science 363, eaau8861 (2019).

28. Davies, B. et al. Altering the Binding Properties of PRDM9 Partially Restores Fertility across the Species Boundary. Molecular Biology and Evolution 38, 5555–5562 (2021).

29. Alleva, B., Brick, K., PraXo, F., Huang, M. & Camerini-Otero, R.D. Cataloging Human PRDM9 Allelic Variation Using Long-Read Sequencing Reveals PRDM9 Population Specificity and Two Distinct Groupings of Related Alleles. Frontiers in Cell and Developmental Biology 9(2021).

30. Berg, I.L. et al. Variants of the protein PRDM9 differentially regulate a set of human meiotic recombination hotspots highly active in African populations. Proceedings of the National Academy of Sciences 108, 12378 (2011).

31. Jeffreys, A.J., CoXon, V.E., Neumann, R. & Lam, K.-W.G. Recombination regulator PRDM9 influences the instability of its own coding sequence in humans. Proceedings of the National Academy of Sciences 110, 600–605 (2013).

32. Paigen, K. & Petkov, P.M. PRDM9 and Its Role in Genetic Recombination. Trends in Genetics 34, 291–300 (2018).

33. Hinch, A.G. et al. The landscape of recombination in African Americans. Nature 476, 170–175 (2011).

34. Buard, J. et al. Diversity of Prdm9 zinc finger array in wild mice unravels new facets of the evolutionary turnover of this coding minisatellite. PloS one 9, e85021–e85021 (2014).

35. Vara, C. et al. PRDM9 Diversity at Fine Geographical Scale Reveals Contrasting Evolutionary PaXerns and Functional Constraints in Natural Populations of House Mice. Molecular Biology and Evolution 36, 1686–1700 (2019).

36. Oliver, P.L. et al. Accelerated Evolution of the Prdm9 Speciation Gene across Diverse Metazoan Taxa. PLOS Genetics 5, e1000753 (2009).

37. Schwartz, J.J., Roach, D.J., Thomas, J.H. & Shendure, J. Primate evolution of the recombination regulator PRDM9. Nature Communications 5, 4370 (2014).

38. Ahlawat, S. et al. Evolutionary dynamics of meiotic recombination hotspots regulator PRDM9 in bovids. Molecular Genetics and Genomics 292, 117–131 (2017).

39. Ahlawat, S., Sharma, P., Sharma, R., Arora, R. & De, S. Zinc Finger Domain of the PRDM9 Gene on Chromosome 1 Exhibits High Diversity in Ruminants but Its Paralog PRDM7 Contains Multiple Disruptive Mutations. PLOS ONE 11, e0156159 (2016).

40. Ahlawat, S. et al. Evidence of positive selection and concerted evolution in the rapidly evolving PRDM9 zinc finger domain in goats and sheep. Animal Genetics 47, 740–751 (2016).

41. Raynaud, M. et al. PRDM9 drives the location and rapid evolution of recombination hotspots in salmonid fish. PLOS Biology 23, e3002950 (2025).

42. Hoge, C. et al. PaXerns of recombination in snakes reveal a tug-of-war between PRDM9 and promoter-like features. Science 383, eadj7026 (2024).

43. Bruno, M., Mahgoub, M. & Macfarlan, T.S. The Arms Race Between KRAB–Zinc Finger Proteins and Endogenous Retroelements and Its Impact on Mammals. Annual Review of Genetics 53, 393–416 (2019).

44. Baker, Z., Przeworski, M. & Sella, G. Down the Penrose stairs, or how selection for fewer recombination hotspots maintains their existence. eLife 12, e83769 (2023).

45. Genestier, A., Duret, L. & Lartillot, N. Bridging the gap between the evolutionary dynamics and the molecular mechanisms of meiosis: A model based exploration of the PRDM9 intra-genomic Red Queen. PLOS Genetics 20, e1011274 (2024).

46. Mihola, O., Trachtulec, Z., Vlcek, C., Schimenti, J.C. & Forejt, J. A Mouse Speciation Gene Encodes a Meiotic Histone H3 Methyltransferase. Science 323, 373 (2009).

47. BhaXacharyya, T. et al. Mechanistic basis of infertility of mouse intersubspecific hybrids. Proceedings of the National Academy of Sciences 110, E468 (2013).

48. Altemose, N. et al. A map of human PRDM9 binding provides evidence for novel behaviors of PRDM9 and other zinc-finger proteins in meiosis. eLife 6, e28383 (2017).

49. Baker, C.L. et al. Multimer Formation Explains Allelic Suppression of PRDM9 Recombination Hotspots. PLOS Genetics 11, e1005512 (2015).

50. PraXo, F. et al. Recombination initiation maps of individual human genomes. Science 346, 1256442 (2014).

51. Grey, C. et al. In vivo binding of PRDM9 reveals interactions with noncanonical genomic sites. Genome Research 27, 580–590 (2017).

52. Patel, A. et al. DNA Conformation Induces Adaptable Binding by Tandem Zinc Finger Proteins. Cell 173, 221–233.e12 (2018).

53. Wolf, G. et al. KRAB-zinc finger protein gene expansion in response to active retrotransposons in the murine lineage. eLife 9, e56337 (2020).

54. Hussin, J. et al. Rare allelic forms of PRDM9 associated with childhood leukemogenesis. Genome Research 23, 419–430 (2013).

55. Sun, Y. et al. Recombination and mutation shape variations in the major histocompatibility complex. Journal of Genetics and Genomics 49, 1151–1161 (2022).

56. Schwarz, T. et al. PRDM9 forms a trimer by interactions within the zinc finger array. Life Science Alliance 2, e201800291 (2019).

57. Baker, C.L. et al. PRDM9 Drives Evolutionary Erosion of Hotspots in Mus musculus through Haplotype-Specific Initiation of Meiotic Recombination. PLOS Genetics 11, e1004916 (2015).

58. Diagouraga, B. et al. PRDM9 Methyltransferase Activity Is Essential for Meiotic DNA Double-Strand Break Formation at Its Binding Sites. Molecular Cell 69, 853–865.e6 (2018).

59. Cosby, R.L. et al. Supplemental Data for Cosby_et_al_2026 [Dataset]. (Zenodo, 2026).

60. Skene, P.J., Henikoff, J.G. & Henikoff, S. Targeted in situ genome-wide profiling with high efficiency for low cell numbers. Nat. Protocols 13, 1006–1019 (2018).

61. PraXo, F. et al. Meiotic recombination mirrors paXerns of germline replication in mice and humans. Cell 184, 4251–4267.e20 (2021).

62. Zhang, Y. et al. Model-based Analysis of ChIP-Seq (MACS). Genome Biology 9, R137 (2008).

63. Quinlan, A.R. & Hall, I.M. BEDTools: a flexible suite of utilities for comparing genomic features. Bioinformatics 26, 841–842 (2010).

64. Gel, B. et al. regioneR: an R/Bioconductor package for the association analysis of genomic regions based on permutation tests. Bioinformatics 32, 289–291 (2016).

65. Heinz, S. et al. Simple combinations of lineage-determining transcription factors prime cis-regulatory elements required for macrophage and B cell identifies. Molecular cell 38, 576–589 (2010).

66. Cosby, R. et al. Cosby_et_al_2020_Supplemental_Data. (Zenodo, 2020).

67. Ernst, J. & Kellis, M. Chromatin-state discovery and genome annotation with ChromHMM. Nature Protocols 12, 2478–2492 (2017).

68. Spence, J.P. & Song, Y.S. Inference and analysis of population-specific fine-scale recombination maps across 26 diverse human populations. Science Advances 5, eaaw9206 (2019).

69. Auton, A. et al. A global reference for human genetic variation. Nature 526, 68–74 (2015).

70. Mihola, O. et al. Histone methyltransferase PRDM9 is not essential for meiosis in male mice. Genome Research (2019).

71. Heddar, A., Fievez, J., Saraeva, R., Benquey, T. & Jouret, G. Heterozygous PRDM9 truncating variant in a patient with primary ovarian insufficiency. Journal of Human Genetics 70, 667–669 (2025).

72. Narasimhan, V.M. et al. Health and population effects of rare gene knockouts in adult humans with related parents. Science 352, 474–477 (2016).

73. Huang, T. et al. The histone modification reader ZCWPW1 links histone methylation to repair of PRDM9-induced meiotic double stand breaks. bioRxiv 14, 836023 (2019).

74. Mahgoub, M. et al. Dual histone methyl reader ZCWPW1 facilitates repair of meiotic double strand breaks in male mice. eLife 9, e53360 (2020).

75. Wells, D. et al. ZCWPW1 is recruited to recombination hotspots by PRDM9 and is essential for meiotic double strand break repair. eLife 9, e53392 (2020).

76. Yuan, S. et al. Dual histone methylation reader ZCWPW2 links histone methylation to initiation of meiotic recombination. bioRxiv, 2026.05.22.727067 (2026).

77. Ruan, T. et al. A novel dual histone mark reader ZCWPW2 regulates meiotic recombination through lactylation and transcriptional regulation in humans and mice. Nucleic Acids Research 54(2026).

78. Schlegel, P.N. et al. Diagnosis and treatment of infertility in men: AUA/ASRM guideline part I. Fertility and Sterility 115, 54–61 (2021).

79. Edgar, R.C. Muscle5: High-accuracy alignment ensembles enable unbiased assessments of sequence homology and phylogeny. Nature Communications 13, 6968 (2022).

80. Dalfovo, D. & Romanel, A. Analysis of Genetic Ancestry from NGS Data Using EthSEQ. Current Protocols 3, e663 (2023).

81. Li, H. Minimap2: pairwise alignment for nucleotide sequences. Bioinformatics 34, 3094–3100 (2018).

82. Li, H. et al. The Sequence Alignment/Map format and SAMtools. Bioinformatics 25, 2078–2079 (2009).

83. Andrews, S. FastQC: A Quality Control Tool for High Throughput Sequence Data. (2010).

84. Martin, M. Cutadapt removes adapter sequences from high-throughput sequencing reads. EMBnet.journal*; Vol* 17*, No* *1*: *Next Genera4on Sequencing Data Analysis* (2011).

85. Langmead, B. & Salzberg, S.L. Fast gapped-read alignment with Bowtie 2. Nature Methods 9, 357–359 (2012).

86. Li, H. & Durbin, R. Fast and accurate short read alignment with Burrows-Wheeler transform. Bioinformatics (Oxford, England) 25, 1754–1760 (2009).

87. Landt, S.G. et al. ChIP-seq guidelines and practices of the ENCODE and modENCODE consortia. Genome Research 22, 1813–1831 (2012).

88. Daley, T. & Smith, A.D. Predicting the molecular complexity of sequencing libraries. Nature Methods 10, 325–327 (2013).

89. Nordin, A., Zambanini, G., Pagella, P. & Cantù, C. The CUT&RUN suspect list of problematic regions of the genome. Genome Biology 24, 185 (2023).

90. Amemiya, H.M., Kundaje, A. & Boyle, A.P. The ENCODE Blacklist: Identification of Problematic Regions of the Genome. Scientific Reports 9, 9354 (2019).

91. Ramírez, F. et al. deepTools2: a next generation web server for deep-sequencing data analysis. Nucleic Acids Research 44, W160–W165 (2016).

92. Halldorsson, B.V. et al. Characterizing mutagenic effects of recombination through a sequence-level genetic map. Science 363, eaau1043 (2019).

93. Spence, J.P. Human Recombination Maps. (Zenodo, 2019).

94. Stovner, E.B. & Sætrom, P. PyRanges: efficient comparison of genomic intervals in Python. Bioinformatics 36, 918–919 (2020).

95. Pedregosa, F. et al. Scikit-learn: Machine Learning in Python. J. Mach. Learn. Res. 12, 2825–2830 (2011).

96. Virtanen, P. et al. SciPy 1.0: fundamental algorithms for scientific computing in Python. Nature Methods 17, 261–272 (2020).

97. Terpilowski, M.A. scikit-posthocs: Pairwise multiple comparison tests in Python. Journal of Open Source SoXware 4(2019).

98. Hunter, J.D. Matplotlib: A 2D Graphics Environment. Computing in Science & Engineering 9, 90–95 (2007).

99. Waskom, M.L. seaborn: statistical data visualization. Journal of Open Source SoXware 6, 3021 (2021).

